# Supramolecular assembly of collagen mimetic peptide D-periodic fibrils and nanoassemblies

**DOI:** 10.1101/2025.02.15.637692

**Authors:** Carson C. Cole, Mark A. B. Kreutzberger, Kevin Klein, Kiana Cahue, Brett H. Pogostin, Adam C. Farsheed, Joseph W.R. Swain, Thi H. Bui, Arghadip Dey, Jonathan Makhoul, Marija Dubackic, Antara Pal, Ulf Olsson, Anđela Šarić, Edward H. Egelman, Jeffrey D. Hartgerink

## Abstract

The collagen triple helix assembles hierarchically into bundled oligomers, solvated networks and fibers. Synthetic peptide assemblies, driven by supramolecular interactions, can form single triple helices through intrahelical amino acid pairs, but the principles guiding interhelical associations into higher-order structures remain unclear. Here, we incorporate cation-*π* and electrostatic charge pairs to probe interhelical interactions and elucidate the mechanisms driving triple helix assembly into fibrils, nanotubes, and nanosheets. Introducing cation-*π* pairs into a fibrillating collagen mimetic resulted in D-periodic fibrils with pH-sensitive gelation. Modifying the presentation of interhelical interactions also enabled the characterization of another D-periodic fibril resembling cartilage collagens, featuring inner and outer triple helix layers. Enhancing electrostatic charge pairs promoted antiparallel assembly, leading to the formation of nanotubes and nanosheets. The packing behavior of triple helices correlates with the interhelical interactions, where parallel associations favor fibril formation, and antiparallel interactions drive nanotube and nanosheet assembly.

## Introduction

Fibrous proteins comprise most of the extracellular matrix (ECM) by mass.^1,2^ Fibronectin, laminin, elastins and collagens are interwoven to provide structural support, template cellular growth and regulate cellular processes.^3,4^ Collagen is classified into over twenty-eight different types and serves as a component for over forty collagen-like proteins.^5^ Fibrous collagens comprise types I, II, III, V and XI.^6^ While the fibrous assembly of the ECM is interdependent, understanding collagen fiber assembly is important due to its role in deleterious diseases^7,8^ The synthesis of collagen-like molecules that recapitulate the hierarchical assembly of collagen also presents an opportunity to Collagen mimetic peptides (CMPs) are relatively short peptide biomimetics that can assemble into collagen triple helices and have served as a model to elucidate the self-assembly of natural collagens. Three monomer strands of left-handed polyproline type II (PPII) secondary structures subsequently nucleate, propagate down the axis and wind into a righthanded supercoiled triple helix.^9,10^ The canonical sequence of these PPII strands is a Xaa-Yaa-Gly repeat, where Xaa is frequently proline (P), and Yaa is 4-hydroxyproline (O). The glycine (G) in the third position is crucial for an axial hydrogen bond and is also sterically favorable for triple helix formation.^11,12^ In addition, the optimized hydrogen bonding network imparted by the sterically unimposing glycine also gives rise to a stagger of a single amino acid between the leading, middle and trailing strands (see Figure 1a) which defines a register between each strand. As shown in Figure 1b, pairwise interactions can occur between adjacent strands in the triple helix in an axial or lateral geometry.^13^ These intrahelical pairwise interactions have been used to control triple helix design by quantifying the relative interaction strength.^14,15^ For example, the electrostatic charge pair of lysine and aspartic acid is stronger than when the cation is arginine. Conversely, cation-π pairs are strongest when arginine is paired with tyrosine rather than lysine in the context of intrahelical interactions.

**Figure 1:**
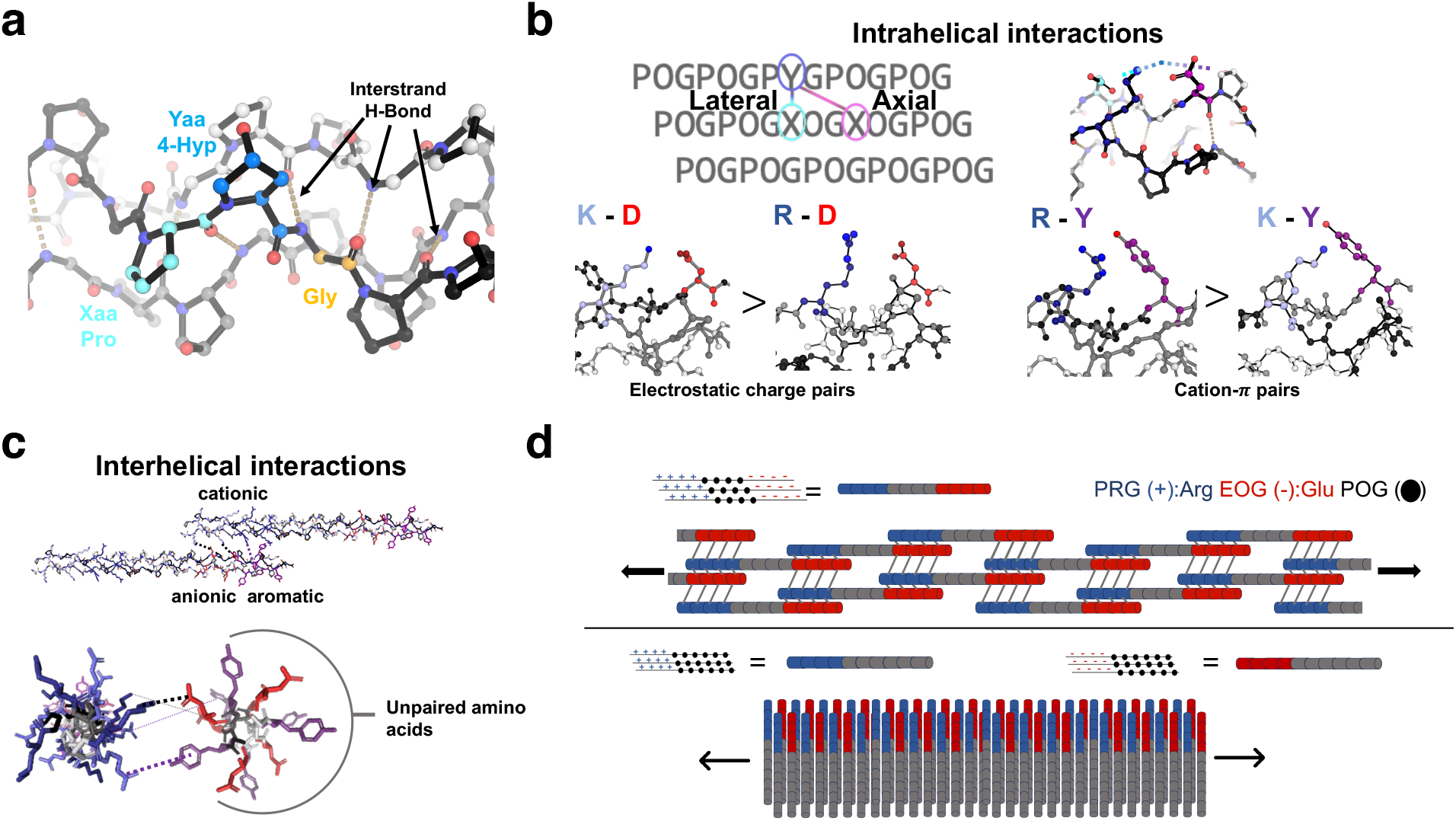
Supramolecular stabilization of collagen mimetic peptides through intrahelical versus interhelical interactions. **a,** The interstrand hydrogen bonding network between the glycine N – H and carbonyl of the amino acid in the Xaa position is key in stabilizing the triple helix. **b,** Intrahelical pairwise amino acid interactions between adjacent strands interact generally in an axial or lateral manner. Axial electrostatic charge pairs and cation-π pairwise interactions have been described with their relative interaction strength and have been used extensively in de novo design. **c,** Proposed models of interhelical electrostatic and cation-π interactions between two triple helices illustrates how interhelical supramolecular interactions are not constrained by the triple helical backbone. **d,** Upper: Staggered association of electrostatic charge pair triple helices have been proposed to assemble into CMP fibrils by interhelical charge pair association in both an axial and lateral growth mechanism.^16^ Lower: A heterogeneous mixture of triple helices has been proposed to associate by electrostatic interactions and assemble laterally into nanotubes and sheets.^17^

Higher-order assembly is driven by interhelical interactions. The atomistic presentation of amino acids that drive self-assembly has yet to be confirmed at an atomistic level until recently for discrete triple helical oligomers but has yet to be extended to the fibril regime.^18,19^ As shown in Figure 1c, associating triple helices possess more degrees of freedom and are not constrained by the backbone of the triple helix. The lack of examples of interhelical associations has limited the development of de novo design rules to design higher-order assemblies. Briefly, self-assembly mechanisms that result in exclusively interhelical interactions have also been proposed to form fibrils and other peptide nanostructures. By utilizing the electrostatic charge pairs of arginine (R) and glutamate (E), a fiber-forming CMP with the sequence (PRG)_4_(POG)_4_(EOG)_4_ was the first synthesized D-Banding CMP fibril. The periodicity was reported to be around 18 nm and hypothesized to assemble with a blunt-ended association mechanism as shown in Fig. 1d.^16^ In addition, the substitution of electrostatic charge pairs has also led to the formation of various collagen mimetic peptide-based assemblies, including nanotubes and nanosheets, which were shown to be pH-dependent.^17,20,21^

Synthesizing fiber-forming CMPs with supramolecular assembly has remained a difficult challenge because of the reliance on passive mechanisms that mediate self-assembly, unlike natural collagen’s active, enzymatically and sterically-guided assembly.^22–25^ However, the self-assembly of higher-order CMPs has been achieved by utilizing supramolecular phenomena such as hydrophobic interactions,^26^ and metals.^27–29^ In addition, aromatic amino acids at the termini of CMPs, such as phenylalanine, have also been reported to assemble CMP fibrils.^30^ Protein expression of a bacterial collagen-like protein flanked with electrostatic interactions has also been shown to generate D-banding fibrils.^31^

The most successful approach to date for designing supramolecular, self-assembling collagen mimetic fibrils has been through employing electrostatic interactions between cationic amino acids, such as lysine (K) or arginine (R), and anionic amino acids, glutamate (E) or aspartate (D). These charge pairs have been proposed to promote assembly by interacting in a pairwise manner intrahelically and interhelically, forming triple helices and then associating into microfibrils as shown in Fig. 2a. In Fig. 2b, a proposed assembly mechanism that utilizes K – D electrostatic interactions has resulted in successful fibrillogenesis^32^ and hydrogel formation by satisfying all theoretical intrahelical pairwise interactions.^33^ The use of electrostatic charge pairs in the peptide (PKG)_4_(POG)_4_(DOG)_4_ (F_0_), as shown in Fig. 2c, led to the first synthetic fibrous collagen mimetic hydrogel.^34^ The fiber design was proposed to interact in a sticky-ended manner, leading to intrahelical interactions and unpaired sets of cationic and anionic regions that could interact interhelically.^35^ As outlined above in Fig. 1d, the blunt-ended assembly of triple helices has also been proposed to associate exclusively by interhelical interactions (see Fig. 2d). However, there has yet to be a comprehensive study that probes how all of these assembly mechanisms are intertwined; here, we present a means of probing such assembly paradigms with cation-π interactions.

**Figure 2:**
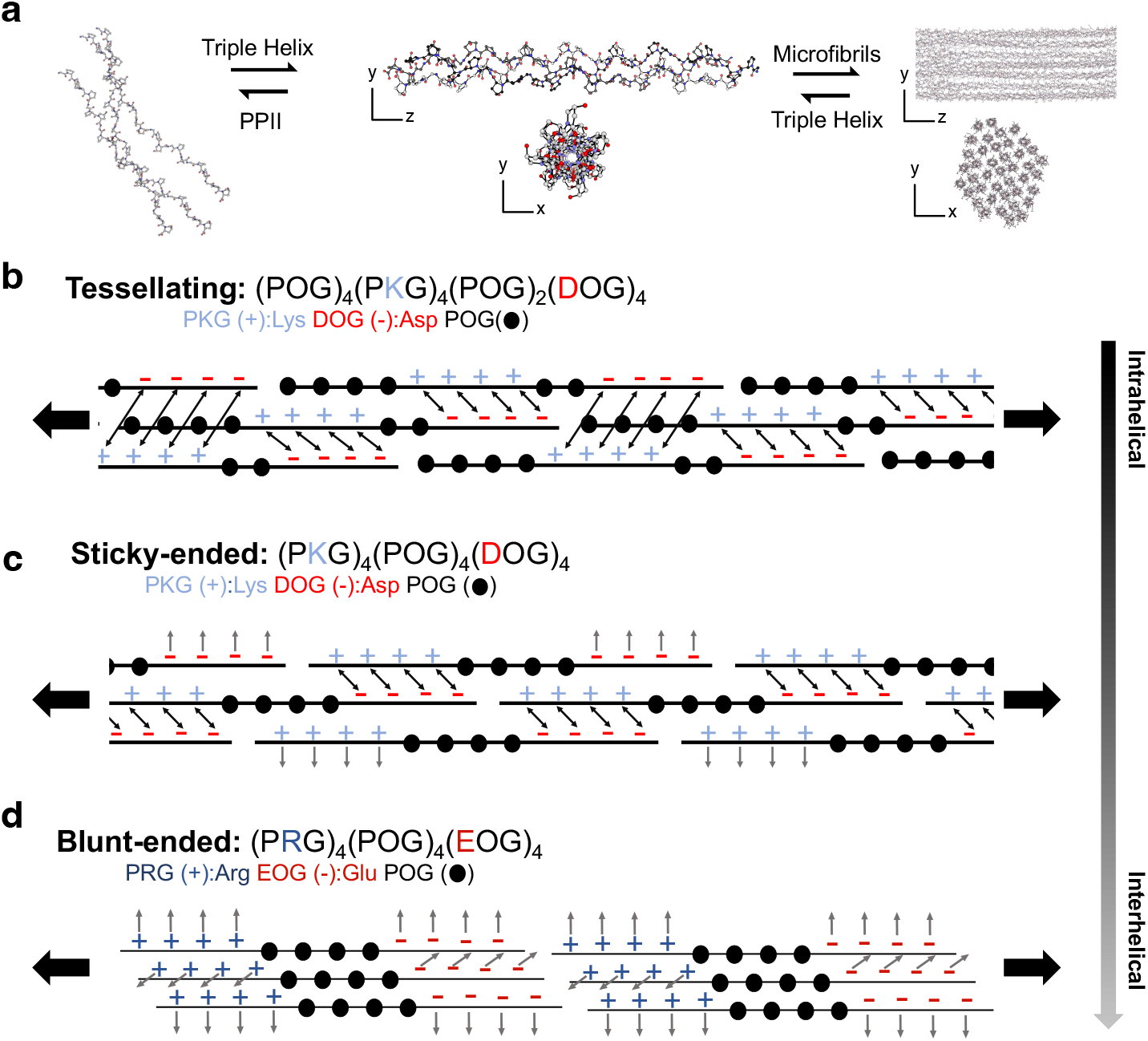
Intrahelical versus interhelical assembly mechanisms of collagen mimetic peptides. **a,** The hierarchical assembly of collagen commences with polyproline type II (PPII) helices assembling into a right-handed triple helix.^36^ These helices then associate into fibrils.^37^ **b,** Tessellation utilizes symmetry rules to pair all theoretically possible interactions between each PPII strand of the triple helix (black arrows).^32^ **c,** Sticky-ended triple helix assembly relies on intrahelical electrostatic interactions between the leading to middle and middle to trailing PPII strands (black arrows). This assembly strategy leaves unpaired sets of electrostatic interactions on the leading and trailing strands (gray arrows) which can interact interhelically.^34^ **d,** Blunt-ended assembly of triple helices has been proposed to assemble through electrostatic interactions. Each triple helix is assembled, and then all available charge pairs form interhelically (gray arrows).^16^

In this work, we optimized polymerizing collagen triple helices can form fibrils through having triplets containing cation-π and electrostatic charge pair interactions. Second, the presentation of theoretically possible pairwise interactions significantly influence solubility and self-assembly dynamics, which can be modulated by adjusting the pH. Additionally, we show that fibrillating collagen mimetics self-assemble from blunt-ended helices that arrange into discrete D-periodic fibrils. These fibrillar structures exhibit triple-helical packing similar to that of natural cartilage collagens. Finally, we demonstrate that these fibrils are structurally related to nanotubes and nanosheets, which can be intentionally designed. These nanoassembles form by promoting a lateral and antiparallel, rather than axial and parallel, association of the triple helices. By applying our rationale of interhelical parallel or antiparallel association, either fibrils or nanoassemblies can be selectively assembled.

## Results and discussion

### Screening Cation-π Interactions for Fibril Assembly

To design polymerizing fibrillar triple helices, we optimized the supramolecular interactions that can lead to triple helix formation and subsequent higher-order assembly. Previous studies investigating intrahelical charge pairs in the context of CMP assembly have pointed to lysine-aspartic acid (K – D) as the ideal charge pair (rendered in (Fig. 3a). Cation-π interactions are known to be important supramolecular interactions in globular proteins and can stabilize triple helices, but the stability granted to the triple helix by these interactions is largely dependent on the conformation and average distance (typically less than 5 Å) between the amino acids. To this end, we screened the relative stability of CMPs after introducing cation-π interactions to the base fiber-forming sequence (PKG)_4_(POG)_4_(DOG)_4_, referred to in the literature as F_0_,^34^ (Fig. 3a) by monitoring the PPII structures with circular dichroism (CD). Liquid chromatography and electron-spray ionization mass spectrometry (ESI-MS) data used to characterize each peptide can be found in the supporting information.

**Figure 3:**
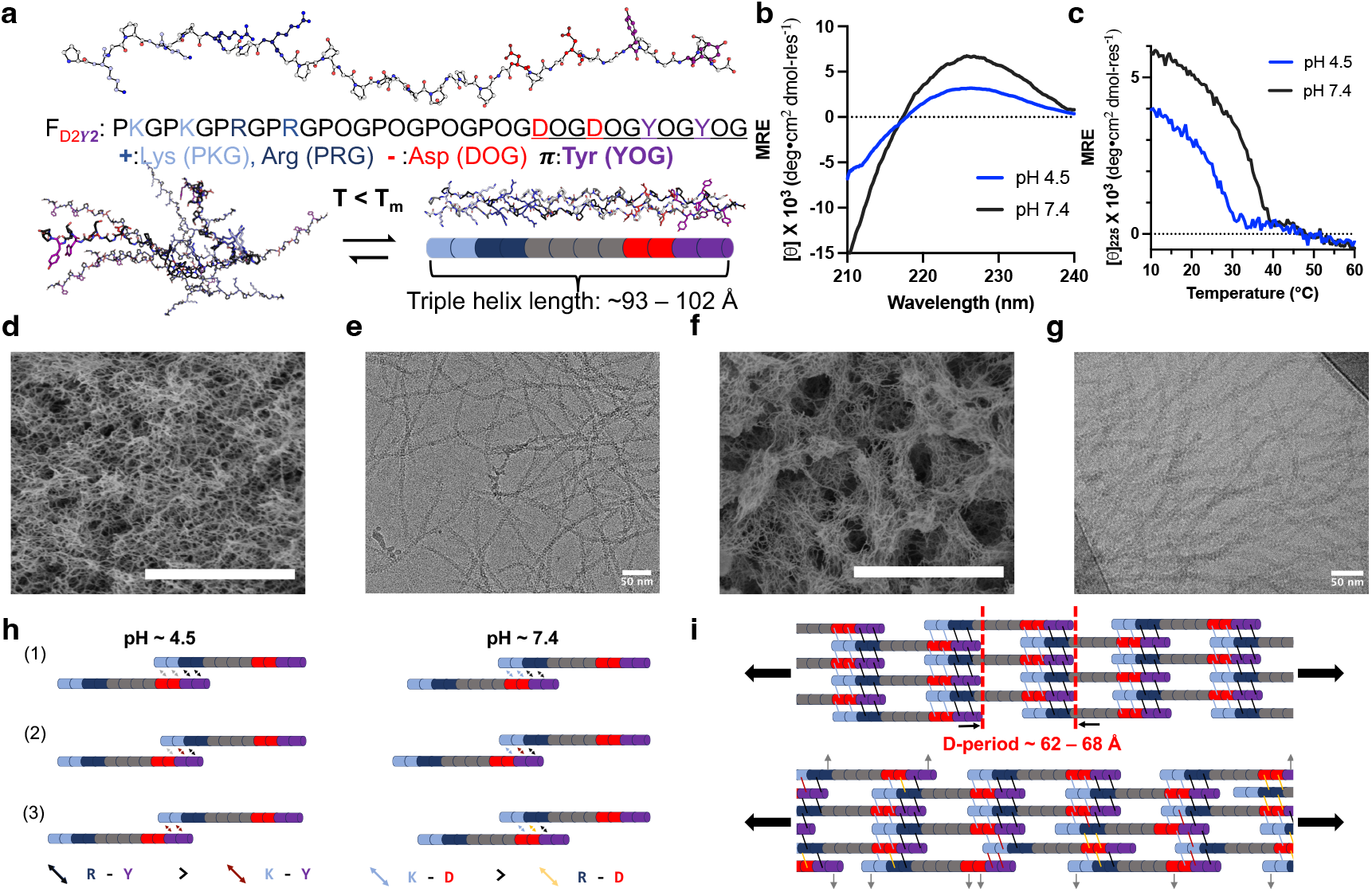
Characterization of F_D2Y2_ supramolecular assemblies at varying pH. **a,** A model of a PPII strand of F_D2Y2_ rendered using AlphaFold3.^38^ Triple helix formation occurs when the temperature (T) is below the *T_m_*. Depending on the winding of the triple helix the length of the helix can range from 93-102 Å. **b,** Circular dichroism (CD) scans shows a strong signal for a PPII helix at a max of *∼*225 nm. **c,** CD Thermal melts reveal pH-dependence of *T_m_* **d,** Scanning electron microscopy (SEM) of F_D2Y2_ at pH 4.5. The scale bar is 3µm. **e,** Cryo-EM of F_D2Y2_, which forms a hydrogel at pH 4.5, shows a fibrous network. The scale bar is 50 nm. **f,** Scanning electron microscopy (SEM) of F_D2Y2_ at pH 7.4. The scale bar is 3µm. **g,** Cryo-EM of F_D2Y2_ shows a clustering of the fibrous assembly as pairwise interaction strengths increase. The scale bar is 50 nm. **h,** Interhelical association of varying cation-π interactions at acidic pH vary compared to neutral pH, where electrostatic charge pairs are enhanced. Relative pairwise interaction strengths are noted. **i,** Fibrillogenesis of F_D2Y2_ shows a periodicity in good agreement with that is observed at pH 4.5. The lower diagram shows how differing staggers can lead to varying periodicities and higher degrees of unpaired interactions that may associate laterally.

For each peptide, F_D2F2_, F_D2W2_ and F_D2Y2_, the PPII signal increased (Fig. 3b-c and supporting information Fig. S8) from acidic to the near neutral pH after adjustment with NaOH and HCl to the specified pH values. All peptides precipitate at pH 9.0 (see Fig. S9). We hypothesize this is due to changes in the protonation states of the charged amino acids and significant changes in solvent interactions upon assembly.^39,40^

All thermal unfolding experimental results are listed in the supporting information Table S1 and the thermal stabilities are defined by the minimum of the derivative (*T_m_*) of each CD folding curve monitored at 225 nm in supporting information Fig. S10. At both pH 4.5 and 7.4, F_D2Y2_ had the highest thermal stability, which increased by 8.0 °C as the pH increased from pH 4.5 to 7.4 (Fig. 3d). With pH adjustment, the stability of F_D2W2_ also increased 12.0 °C and F_D2F2_ increased 6.0 °C. As the pH is adjusted to 7.4, the lysine – aspartate interactions contribute more to the thermal stability of the triple helix than at pH 4.5. With this assumption, it is clear that the aromatic groups show the same destabilizing effect in the fibrous systems as they do in discrete triple helices.^41^ We selected the tyrosine-containing F_D2Y2_ assembly for further investigation by Cryo-EM due to the high thermal stability and gel-like composition of this peptide at pH 4.5.

### A comparison of intrahelical versus interhelical fibril assembly

Collagen mimetic peptide fibrillogenesis, like natural collagen, is dependent on a variety of factors in the aqueous environment,^42,43^ including concentration,^34^ ionic strength,^32^ and pH.^44^ This was also the case for F_D2Y2_. We found that at a concentration of 2% w/v and in MQ water, F_D2Y2_ was soluble at pH 4.5 and formed fibrils. The peptide began to form thicker fibers as the pH increased, leading to slight precipitation at neutral pH. At pH 4.5, the fibrous nature of F_D2Y2_ was confirmed with scanning electron microscopy (SEM) (Fig. 3d) and cryo-EM demonstrated D-periodic fibrils with a periodicity of *∼*55-60 Å as presented in Fig. 3e. F_D2Y2_ formed a hydrogel at acidic pH that exhibited linear elasticity to 8% strain. The storage modulus in the linear viscoelastic range is 39 Pa, and the loss modulus is 9 Pa (Fig. S11). At lower concentrations and in phosphate buffer gelation was not observed.

Characterization for F_D2Y2_ at pH 7.4 in Fig. 3f/g and 3h showed that fibril diameter increased, but D-banding was still observed. Additionally, when dissolved in 10 mM phosphate buffer, F_D2Y2_’s fibril diameter was observed to thicken and the firbil length shorten from 40 nm to 20 nm, as confirmed with both Cryo-EM and small-angle X-ray scattering experiments (Fig. S12), which has previously been observed in the literature.^45^ The triple helices at pH 7.4 exhibit higher thermal stability (Fig. 3d) while still undergoing fibrillogenesis in MQ H_2_O as confirmed with Cryo-EM (Fig. S13 A/B).

The observation of D-periodic fibers suggests that the triple helices of F_D2Y2_ are interacting interhelically and in a blunt-ended manner as proposed in Fig. 2d. In a simplistic model of interhelical interactions, we hypothesize that the staggers of triple helices can give rise to different pairwise interactions. The three most stable interhelical staggers are presented in Fig. 3h, and a diagram of all possible parallel staggers is included in the supporting information (Fig. S14). Cation-π pairs are assumed to be stronger when arginine, not lysine, is the cation,^46^ and electrostatic charge pairs usually favors lysine to pair with aspartate.^47^ At acidic pH, we assume that the electrostatic interactions do not contribute as significantly to interhelical association compared to the cation-π pairs, which influences in the discrepancy in associations that were observed with Cryo-EM.

Triple helices are proposed to have roughly a 10/3V or 7/2V symmetry, depending on the density of proline-rich amino acid residues.^11,36^ This symmetry defines the axial height of a helix and rise for each triplet in the sequence. For a 10/3V triple helix, the rise of each triplet is 7.8 Å, which would roughly yield a 102 Å height for the 36 residues in F_D2Y2_’s sequence. If calculated using the 7/2V model, with a rise of 8.57 Å per triplet, the helix would be roughly 94 Å. As illustrated in Fig. 3i, this would result in a periodicity ranging from 62 – 68 Å, which is close to the 55-60 Å periodicity observed from Cryo-EM. This periodicity assumes all pairwise interactions were in their most stable pairing, but it would also be reasonable for the helices to stagger leading to different periodicity and unpaired interactions (rendered in gray, Fig. 3i) which could drive lateral packing and thickening of fibrils.

We then compared the fibrous CMP of F_D2Y2_ with the previously established F_0_,^34^ see Fig. 2c, and a tessellating collagen mimetic peptide with cation-π interactions (F_D2Y2_*_−_*_T_) to compare fiber morphology. Analysis of F_0_, which is proposed to assemble with both intrahelical and interhelical electrostatic charge pairs, at pH 7.4 in 10 mM phosphate buffer resulted in fibrils that possessed distinct morphological differences compared to F_D2Y2_. Circular dichroism showed asymmetry in the thermal unfolding curves but an overall gain in thermal stability as the pH increased (Fig. S15a-c). At pH 4.5, the sample did not assemble into a hydrogel, supporting that the electrostatic charge pairs are weaker near the pKa of aspartic acid. The sample formed a hydrogel at 1.0 % w/v and pH 7.4, as previously reported. Cryo-EM of this hydrogel revealed fibrils of varying diameters and lacking periodicity (Fig. S15d), and SEM also supports the distinctness of F_0_’s assembly when compared to F_D2Y2_ (Fig. S15e/f).

Next, we used CD and cryo-EM to analyze F_D2Y2_*_−_*_T_. Fig. S16a of the supporting information shows the proposed assembly mechanism of F_D2Y2_*_−_*_T_ and the cation-π substitution scheme into the 42mer peptide. The main feature of this assembly paradigm is that there are theoretically no interhelical interactions. The assembly of F_D2Y2_*_−_*_T_ appears distinct from F_D2Y2_ and F_0_ in the fact that the adjustment of pH, from 4.5 to 7.4 in MQ H_2_O, did not substantially increase the thermal stability (Fig. S16b-d). We also observed with Cryo-EM (Fig. S16e/f) that at pH 4.5 there were two distinct assemblies: fibrous and aggregated. The fibrils were notably smaller in diameter compared to F_D2Y2_ and F_0_. We hypothesize that the aggregation occurs because tyrosine is sterically larger than aspartic acid and may disrupt axial, long-range assembly. These data suggest that the resulting fibrils from F_0_, F_D2Y2_*_−_*_T_ and F_D2Y2_ are distinct collagen mimetic peptide assemblies.

### Enhanced D-Periodicity by alternating interaction pairs

Subtle changes to amino acid sequences can change the structure of peptide supramolecular assembly.^48–50^ To probe how alternating triplets of each pairwise interaction can impact assembly, we designed an alternating fiber-forming peptide termed F_DY2_. In this design, the termini alternate between each type of pairwise interaction (Fig. 4a). While the parallel association leads to two cation-π pairs and two electrostatic charge pairs, the competing interhelical staggers would be distinct from F_D2Y2_. The CD thermal melt of the two samples, shows a distinct *T_m_* between the samples (Fig. 4b). In addition, as reported in supporting information Table S2, the samples also precipitated at different rates, with F_DY2_ being insoluble at pH 7.4. However, F_DY2_ also formed a relatively weak hydrogel at pH 4.5 in MQ H_2_O, and the loss and storage moduli are approximately 32 Pa and 5 Pa in the linear regime, respectively (see Fig. S12).

**Figure 4:**
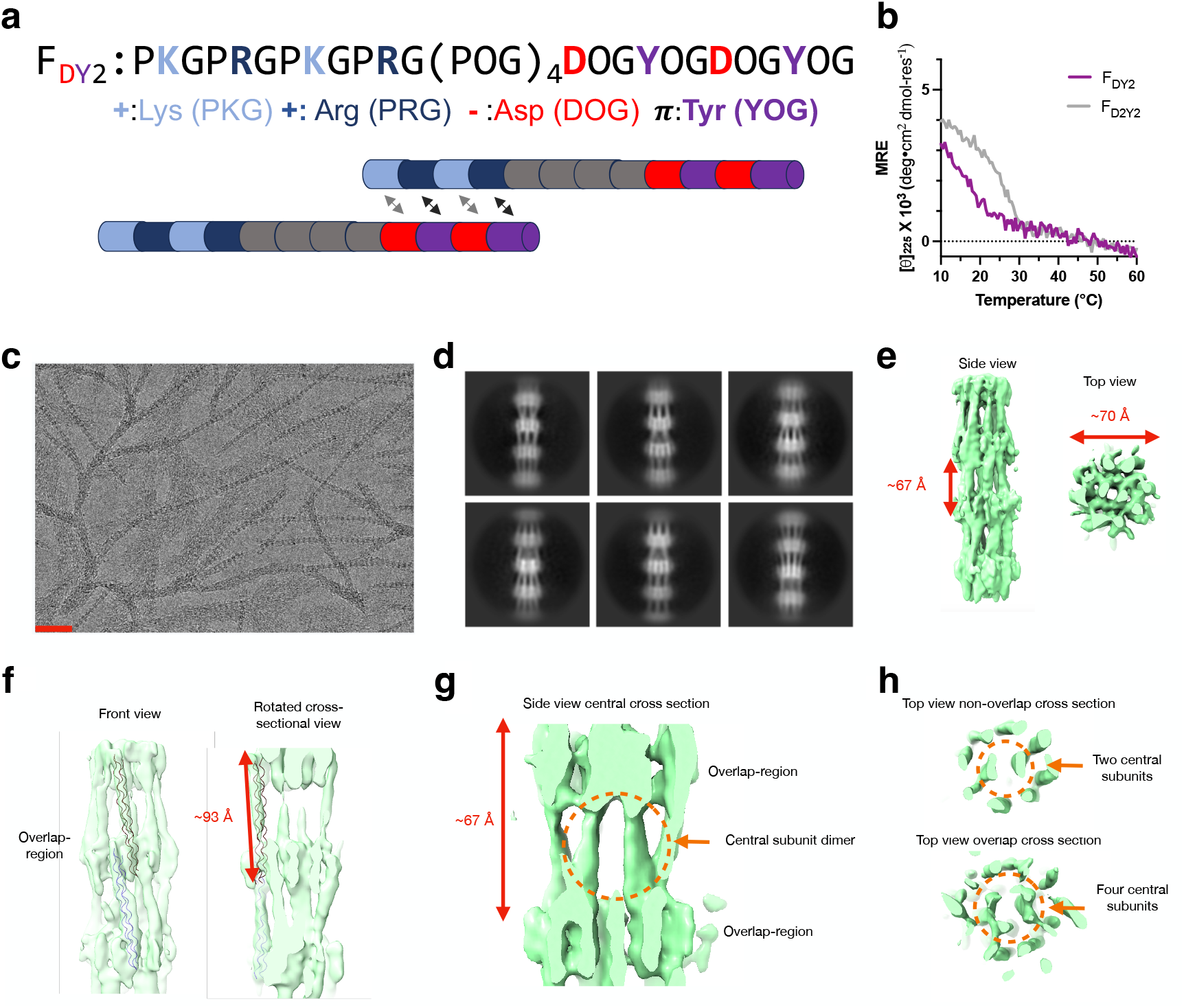
Alternating triplets of pairwise interactions in F_DY2_ yield triple helical D-banding. **a,** Sequence and the most stable parallel interaction between two triple helices of F_DY2_. **b,** Thermal melting curves of F_DY2_ compared with F_D2Y2_ yield different thermal stabilities. **c,** A representative Cryo-EM micrograph of F_DY2_ with fibrous morphology that possess D-banding periodicity. The scale bar is 50 nm. **d,** 2D class averages of the F_DY2_ fibrils revealing a D-banding pattern. **e,** 3D reconstruction of particles selected from the best 2D class averages showing a *∼*67 Å period and *∼*70 Å diameter. **f,** Using a collagen mimetic peptide crystal structure (PDB ID: 6JEC), the 3D volume could be correlated to triple helices winding together. **g,** A lateral slice of the 3D volume shows a central dimer unit in the nonoverlapping region. **h,** Axial slices compare the cores of the non-overlapping regions showing the two central helices, as opposed to the overlapping region with four central subunits. The overlapping region is hypothesized to be where the electrostatic charge pairs and cation-π pairwise interactions occur. All data were collected at pH 4.5.

We further investigated how alternating interactions would impact the fibril formation of F_DY2_ with Cryo-EM. Micrographs in Fig. 4c showed distinct fibrils with D-banding characteristics, a hallmark of natural collagen. Further processing of the cryo-EM data resulted in 2D class averages (Fig. 4d) with obvious periodicity and possibly showed individual triple helices. Then, the best 2D classes with similar diameters were chosen for subsequent 3D reconstruction, resulting in the density map shown in Fig. 4e. We measured that fibrils had a 67 Å periodicity, which is 1/10 of the 670 Å periodicity of natural human collagen type I.^51,52^

Also apparent in this 8 Å resolved density map were rod-like densities, which matches the expected shape and morphology of a collagen triple helix at that resolution (Fig. S17). Using a deposited crystal structure of a triple helix (PDB ID: 6JEC), we fit a triple helix corresponding to a length of 93 Å into the putative low-resolution triple helix density in Fig. 4f. Fitting a second model into the density showed how these triple helices might overlap. The periodicity of 67 Å is nearly 2/3 the length of the helix, which could encompass either the cationic/POG sequence (PKGPRG)_2_(POG)_4_ triplets or the anionic and aromatic/POG regions (POG)_4_(DOGYOG)_2_. This would lead to the last or first four triplets being unpaired, which could interact with the next set of helices. This is similar to the previously reported parallel blunt-ended assembly by Rele.^16^

We found that this was further supported by the apparent arrangement of non-overlapping regions, which appeared to have a central dimer of triple helices surrounded by eight triple helices (Fig. 4g/h), termed an 8+2 structure. At the overlapping region, four central subunits are possibly surrounded by sixteen outer triple helices in a 16+4 arrangement (Fig. 4f). Models are proposed in supporting information in Fig. S18. This is also observed in cartilage fibrils, where the triple helices are hypothesized to be arranged in a 10+4 bundling structure.^53^ These data also demonstrate that supramolecular assembly can be used to create collagen mimetics that are relatively uniform in their lateral packing and may be used to better understand how lateral packing is curtailed in natural collagens.^22,54^

### Formation of collagen mimetic nanotubes and nanosheets by antiparallel assembly

To further probe how pairwise interactions affect self-assembly, we synthesized F_YD2_. This amino acid sequence, presented in Table S1, alternates pairwise interactions like F_DY2_, but upon parallel association should favor the weaker R – D electrostatic charge pair and K – Y cation-π pair by flipping only the anionic and aromatic region of the sequence but keeping the cationic region constant. Upon dilution in MQ H_2_O and pH adjustment, we observed precipitate formation for each solution at pH 4.5, 7.4 and 9.0 (Fig. S19), which suggested a different mechanism of association.

An antiparallel association of triple helices can lead to the formation of nanotubes and nanosheets as shown in Fig. 5a and has been previously reported to be formed with a similar 36mer peptide.^17,21^ However, a fibrous morphology leading to this transition has yet to be reported. The hypothesized configurations of the three most stable staggers of F_DY2_ and F_YD2_ are presented in Fig. 5b. At pH 7.4 we consider electrostatic interactions to have a significant influence on F_DY2_ changing from fibrils to nanotubes due to the stabilizing, antiparallel staggers in (2) and (3). Cryo-EM experimental data presented in Fig. 5c confirmed nanotube formation. Additional supporting information data show that the tubes can also be multi-lamellar (Fig. S20a/b). Conversely, F_YD2_ has the highest degree of satisfied interactions in the antiparallel stagger of (1). We confirmed with Cryo-EM that this peptide formed nanosheets at pH 7.4. These nanosheets were substantially larger both the x and y dimensions than the tubes but also appear to have some curling at the edges resulting in darker edges in the micrograph in Fig. 5d. Fig. S20c/d demonstrates more cases of larger nanosheets and a side profile of the curling. In addition, CD of the supernatant of these solutions showed that triple helices had a higher thermal stability as the pH increased (Fig. S21).

**Figure 5:**
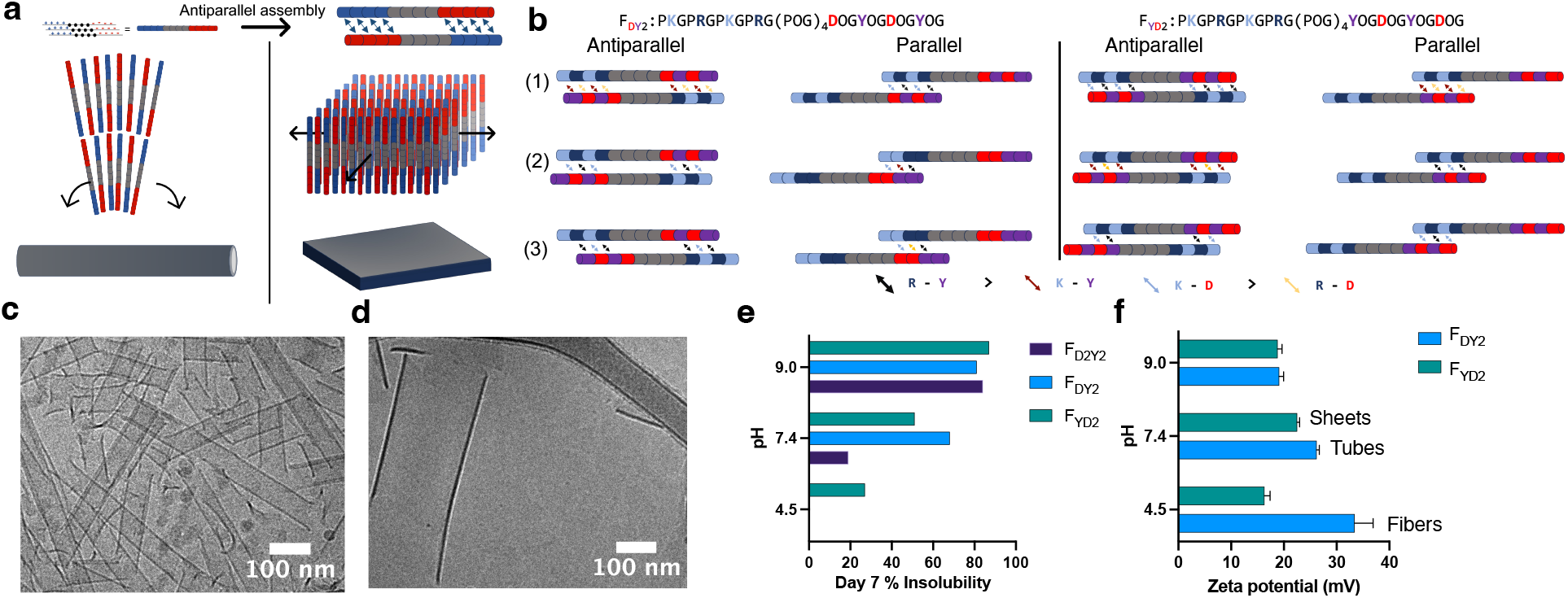
Antiparallel association of triple helices leads to nanotube and nanosheets at neutral pH. **a,** Triple helices can also interact in an antiparallel manner, which satisfies a set of pairwise interactions at both the N and C-termini. This association is hypothesized to drive tubular or sheet-like structures. **b,** Comparison of F_DY2_ and F_YD2_’s propensity to associate in a parallel or antiparallel manner with the three most stable shifts for each association paradigm. **c,** Cryo-EM of F_DY2_ shows unilamellar and multi-lamellar nanotubes at pH 7.4. **d,** Representative Cryo-EM of F_DY2_ at pH 7.4 shows nanosheet formation. Scale bars are 100 nm. **e,** A bar graph of 2.0 wt % (v/v) samples of F_DY2_ and F_YD2_ after seven days shows pH-dependent rates of precipitation. **f,** A bar graph of F_DY2_ andF_YD2_ zeta potentials at varying pH demonstrates a difference in overall charge of the microfibrils, nanotubes and nanosheets.

Using the tyrosine tags of F_YD2_, F_DY2_ and F_D2Y2_, we performed a mass balance of each peptide due to each sample, despite having a remarkably similar amino acid composition, exhibiting different solubilites in water. F_D2Y2_ had the highest solubility at pH 4.5, 7.4 and 9.0, possibly due to the staggering as proposed above. The results, summarized in Table S2 show that F_YD2_ exhibit significantly lower solubility than F_DY2_. In Fig. 5d the comparison of F_YD2_ and F_DY2_ compares the disparity between the precipitation rate of the two peptides after a week. We performed zeta potential measurements, see Fig. 5e, observing that fibers carry excess positive charge. This could help explain the thickening of fibrils upon introduction of salt-containing buffers, where longer-range parallel association is disrupted. Nanotube/nanosheet zeta potentials suggest similar charge at neutral and basic pH values.

To further investigate the nuances of parallel association between F_D2Y2_ and F_DY2_ leading to fibrils, we used a coarse-grained model. Coarse grained molecular dynamics (MD) simulations have successfully been used in understanding how amino acid based intermolecular interactions guide and influence self-assembly of collagen and collagen mimetic fibrils.^43,55,56^ To understand how different combinations of peptide designs and pH conditions influence the assembly process, we carried out iterative fibrillogenesis MD simulations for F_DY2_, F_D2Y2_and F_YD2_. In our MD simulations, the CMPs are modeled as flexible rods that carry the physical properties of the CMP amino acid sequence (Fig. 6a). Two such rods can mutually interact via electrostatic as well as cation-π pairwise interactions. Both kinds of interactions are modeled using two variations of a Debye-Hückel potential. To mimic a change in pH, we vary the strength of the electrostatic interactions while leaving the cation-π interactions unchanged.

**Figure 6:**
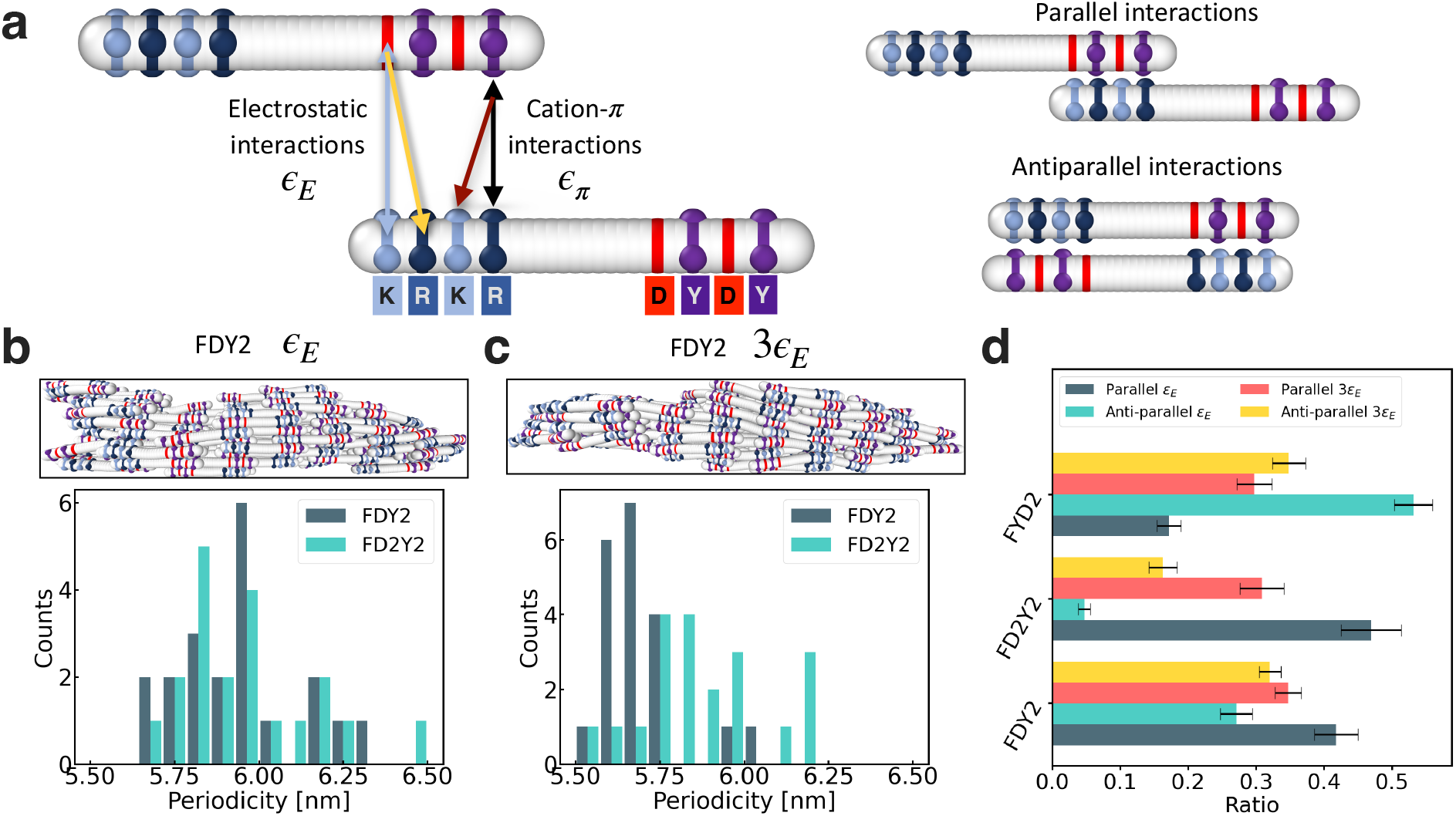
Coarse-grained simulations of periodicity and triple helix association. **a,** Coarsegrained flexible rods corresponding to a blunt-ended triple helix of F_DY2_. Light blue strips represent lysine, dark blue arginine, red aspartic acid and purple tyrosine. Relative interactions are denoted with ε. To the right, flexible rods can associate in antiparallel and parallel geometries. **b,** F_DY2_ and F_D2Y2_ fibril formation and measured periodicity with lower electrostatic interaction strength, ε_E_. A snapshot of a representative F_DY2_ fibril from a simulation. **c,** F_DY2_ and F_D2Y2_ fibril formation and measured periodicity with higher electrostatic interaction strength, 3ε_E_. A snapshot of a representative F_DY2_ fibril from a simulation. **d,** Ratios of parallel versus antiparallel association for F_YD2_, F_D2Y2_ and F_DY2_ rods at lower and higher electrostatic interactions strengths. Error bars represent standard error of the mean (SEM) for each data set for n = 20 simulations.

To study differences in the assembled fibril structure, we ran 20 independent fibrillogenesis simulations for each of the three CMPs at two different electrostatic interactions strengths (ε_E_ and 3ε_E_) The enhancing of electrostatic forces in the 3ε_E_ experiments is meant to replicate the pH adjustment from acidic to neutral pH in our experiments. Snapshots of such simulations are shown in Fig. 6b. We then measured the resulting fibrils for periodicity as well as for their composition of parallel and antiparallel stacking. The histograms of the measured periodicities in Fig. 6c do not reveal substantial differences between F_DY2_ and F_D2Y2_. F_DY2_ has mean periodicity of 5.69 (0.03) nm and 5.95 (0.04) nm at strong and weak electrostatic interactions, respectively, while F_D2Y2_ has mean periodicity of 5.87 (0.04) nm and 5.96 (0.04) nm at strong and weak electrostatic interactions, respectively, where the number in brackets is the standard error of the mean. The only noticeable change is the shift of the mean to larger periodicity for F_DY2_ as the electrostatic interaction strength is reduced. However, when looking at fibril compositions, we can see significant differences between the three CMPs (Fig. 6d).

At strong electrostatic interaction strengths (3ε), we can see that both F_DY2_ and F_YD2_ have only a small gap between antiparallel and parallel stacking. This indicates a larger tendency to be in antiparallel configurations than for F_D2Y2_, which would favor a sheet or tube conformation over a fibril conformation. Meanwhile, F_D2Y2_ shows a large gap already at strong electrostatic interactions, which is in line with the observation that F_D2Y2_ is forming fibrils at strong electrostatic interactions. When we decrease the electrostatic interaction strength, this gap becomes even larger, promoting a periodic fibrillar conformation even further. At the same time, F_DY2_ increases its tendency for parallel configurations while the tendency for antiparallel configurations decreases. This is in agreement with the observation that F_DY2_ starts to form fibrils at lower electrostatic interactions. F_YD2_ instead shows the opposite trend, increasing tendency for antiparallel configurations and decreasing tendency for parallel configurations, further promoting non-fibrillar conformations. This change in fibril composition is in good agreement with the experimental observation and supports the hypothesis that slight changes in amino acid sequences can significantly change the self-assembled macroscopic structure. We are, however, not able to capture the difference in periodicity observed in the experiments or the formation of sheets/tubes for F_DY2_ and F_YD2_. Since our coarse grained model is a simplification of the real triple helical structure, we cannot expect to capture the full self-assembly process in detail nor the register of the triple helices which is crucial in forming supramolecular bonds in PPII structures.^57^ Furthermore, our fibrillogenesis simulation approach is designed to promote the growth of a single fibril and therefore not suited to capture the formation of sheet or nanotube like conformations. Nevertheless, we are able to show how tendencies for parallel vs antiparallel configurations are impacted by changes in the electrostatic interaction strength and presentation of supramolecular interactions. Since fibril and sheet formation is connected to parallel or antiparallel configurations, our model can qualitatively provide an explanation in showing how changes in electrostatic interaction strength lead to different tendencies towards parallel and antiparallel configurations in CMP designs.

Understanding the interactions that give rise to different CMP nanostructures could help guide the de novo design of collagen mimetic nanomaterials with possible applications in bioengineering or medicine. Further work into establishing the atomistic nuances of self-assembly will also inform how to enhance colloidal and thermal stability of these biomimetic materials. Fibrillogenesis is a complex, higher-order problem in assembly that will guide further discoveries on how ECM proteins maintain homeostasis and achieve the correct fiber dimensions. Understanding these processes could help mitigate deleterious pathways such as fibrosis or oncogenesis.^58^ Further incorporation of other supramolecular interactions, and naturally expressed non-canonical amino acids, could also be pursued to understand their role in fibrillogenesis.

## Conclusion

Fibrous collagen mimetic peptides and their design have classically used charge pair interactions to drive higher-order assembly. Here, we demonstrate that a cation-π pairwise interaction can be successfully incorporated into various sequences and is used to differentiate different mechanisms for polymerization. Tyrosine was the best aromatic amino acid to pair with arginine, consistent with previous findings, and also resulted in pH-dependent assemblies. At acidic pH, tyrosine-containing peptides formed a collagen mimetic hydrogel.

Characterization of this peptide at increasing pH showed a thickening of fibrils as supramolecular attractive forces were strengthened, and eventually, the sample precipitated. Finally, by investigating alternating electrostatic charge pair and cation-π containing triplets, we enhanced D-banding periodicity, a hallmark feature of natural collagen. These molecules assemble in a parallel association mechanism, and much like cartilage collagens, possess an inner and outer-layer of triple helices. At neutral pH, these alternating sequences formed nanotubes by antiparallel association. We then showed that the nanotubes could be unfurled into nanosheets by flipping the triplet pairs and favoring anti-parallel associations. Further fundamental studies of the atomistic rationale of these supramolecular assembly mechanisms serve as a basis for the bottom-up design of collagen mimetic biomaterials.

## Experimental

### Circular dichroism

Circular dichroism (CD) spectroscopy was performed on a Jasco J-810 spectropolarimeter (Tokyo, Japan) with a Peltier temperature-controlled stage. Peptide samples were prepared at a 3 mM working concentration and diluted with pH-adjusted MQ water to 0.3 mM for melts. After preparation, 200 µL of the sample was transferred to a quartz cuvette of 0.1 cm path length. Wavelength scans between 180 and 250 nm confirmed the PPII secondary structure. The maximum, which falls near 225 nm, was then monitored with a thermal ramp from 5 to 65 °C at 10 °C/h. Using a first-degree Savitzky–Golay smoothing algorithm, the first derivative curve was calculated for melting curves. The minimum of the first derivative is defined as the melting temperature (T*_m_*). The MRE was calculated as previously reported.^14^

### Scanning electron microscopy

Samples were dehydrated in a series of ethanol:water mixtures (30%, 50%, 60%, 70%, 80%, 90%, 2 × 100% (v/v)). Samples were placed in each bath for 10 min to dehydrate the peptide samples and then further dried using a Leica EM CPD300 Critical Point Dryer (Leica Biosystems, Deer Park, IL). Dried samples were then sputtered with 10 nm of gold to enhance conductivity with a Denton Desk V Sputter System (Denton Vacuum, Moorestown, NJ). Sputtered samples were then imaged on a Helios NanoLab 660 Scanning Electron Microscope (FEI Company, Hillsboro, OR) at 1 kV and 25 pA.

### Cryo Electron Microscopy

Fibrous samples were added to Lacey Carbon CU 300 mesh grids and were frozen with a VitrobotTM Mark IV plunge freezer. Prior to freezing, the grids were subject to Glow discharge treatment using an Electron Microscopy Sciences GloQube plasma cleaner system. Grids were imaged on a Titan Krios equipped with a K3 direct electron detector at 105,000x magnification (0.82 As /pixel). Cryo-EM reconstruction and image processing of raw movies and subsequent image processing and reconstruction steps were handled in cryoSPARC.^59^

For structural determination of the F_DY2_ filaments, helical and single-particle reconstruction was attempted separately. For the single-particle reconstruction, particles belonging to good 2D classes of filaments, were subject to ab initio reconstruction where a volume with the 67 Å periodicity was apparent. The ab initio volume and its corresponding particles were input into a homogeneous refinement job with a mask created from the ab initio. The structure had an apparent C2 symmetry, so a subsequent homogenous refinement job imposing this point group symmetry was run. The resolution of this volume was rather low and to remove additional noise were low-pass filtered to 8 Å.

Even at this low resolution, density was observed for triple helices. While individual chains of the triple helices were not resolvable, existing triple helix crystal structures could esaily fit into the density map. The same particles from the good 2D class averages were used in a helical refinement job using a cylindrical starting volume for helical reconstruction. No final helical volume could be generated that was superior to the volume obtained in ab initio reconstruction. All volumes and figures were analyzed and made in UCSF ChimeraX.^60^ The final cryo-EM volume was deposited with the electron microscopy data bank (EMDB) as EMD-48818.

### Coarse-grained molecular dynamics simulations

The coarse-grained model was adapted from the "D-mimetic" modelvpreviously used to study self-assembly properties.^43,55,56^ The coarse grained versions of the CMPs were modeled as linear chains consisting of 36 beads. Each bead measures a diameter of *D* = 1.12σ, where σ is the MD unit of length. Beads within one molecule are connected to their direct neighbors via a harmonic bond *V* = κ_bond_(*r* *-* *r*_0_)^2^, where *r*_0_ = 0.24σ is the equilibrium length and κ_bond_ = 500kT/σ^2^ controls the bond strength. The CMPs span a length between 9.3 - 10.2 nm and the coarse grained molecules have a length of 9.76σ, thus σ = 1 nm. To model molecular rigidity of the coarse grained molecules, a harmonic angular potential V = κ_angle_(θ *–*θ_0_)^2^ was used. This potential controls the angular movement between three neighboring beads, where θ_0_ = π is the equilibrium angle and κ_angle_ = 500 kT defines the strength of the potential.

Furthermore, inter-molecular interactions were modeled between any two molecules that come into defined interaction ranges. Beads that represented the amino acids lysine (K) and arginine (R) carry a positive dimensionless MD charge Q_K_ = 16.5 and Q_R_ = 3.0, while beads that represented aspartic acid (D) carry a negative dimensionless MD charge Q_D_ = *-*3.375. If any of these three type of beads from two different molecules get close to each other, an interaction occurs via a screened electrostatic potential (DLVO) 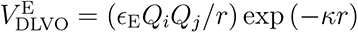, if the distance between the two beads 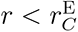 was smaller than the cutoff distance 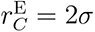. If 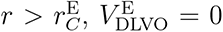 was the Debye screening length and ε_E_ defined the effective interaction strength that was varied.

To model cation-π pairwise interactions, three patches each for the amino acids lysine, arginine, and tyrosine (Y) were introduced. These patches were rigidly connected to their carrying bead at a radial distance of *r*_patch_ = 0.46σ. Additionally, the patches were arranged at rigid angles of 2π/3 between each other to mimic the geometry of the triple helical arrangement. The plane spanned by the three patches was perpendicular to the molecule axis, such that the three patches were oriented towards the surface of the coarse grained molecule. The lysine and arginine patches each carried a dimensionless MD charge *P*_K_ = 2.75 and *P*_R_ = 4.5, while the tyrosine patches each carried a dimensionless MD charge P_Y_ = *-*0.75. Different to the electrostatic pairwise interactions, for the cation-π pairwise interactions only lysine-tyrosine and arginine-tyrosine contacts interact with each other. If the distance between any two patches of those two pairs becomes smaller than the cutoff distance 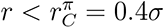, they start interacting via another screened electrostatic potential 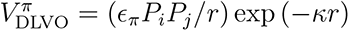, where ε*_π_* is the effective strength of the cation-π interaction. Charges and patches within the same molecule do not interact with each other. At the start of each simulation, the positions and orientations of N = 99 coarse grained molecules were randomized and placed in a cubic volume of dimension *L* = 44σ.

The simulation itself was started in a periodic simulation box of dimension 2L = 88σ, such that the initially placed molecules were centered in the larger simulation volume. Enlarging the simulation volume for the initial fibril nucleation was proven to yield more robust results, as a smaller simulation volume can lead to problems such as fibrils infinitely looping through the periodic boundaries. Using the LAMMPS MD package, the simulation is was ran at constant number of particles N and constant volume V using a Langevin thermostat to mimic Brownian motion of the coarse grained molecules.^61^ With *τ*_0_ being the MD unit of time, the simulation was ran for 10^6^ integration time-steps of size 0.001*τ*_0_ while using a damping coefficient *γ* = 1*τ*_0_. After this initial fibril nucleation simulation, equilibration and growth simulations were alternated. Molecules were provided sufficient time to equilibrate in each simulation. When transitioning from equilibration to growth phase, the largest cluster of molecules that existed at the end of the previous equilibration phase was identified. This cluster was then aligned to the x-axis and re-centered in the simulation box. Then an amount of new randomly oriented coarse-grained monomers N_added_ were added and the box was then resized such that the number concentration of total molecules in the box is c_mol_ = 0.001σ*^−^*^3^. Using this technique, new monomers were iteratively added to the simulation and promoted the growth of a single large fibril.

### Zeta potential measurements

The ζ-potential values for F_DY2_ and F_YD2_ were acquired using a Malvern Zen 3600 Zetasizer (Malvern Instruments Ltd., Malvern, U.K.). Peptide solutions were left to assemble after annealing at 2% (w/v) in MQ water and then were diluted 1:20 in 3 mM NaCl for a final concentration of 0.1 wt% (v/v) and immediately added to a Malvern Zetasizer Nano Series folded capillary cell. Each sample was measured three times and then averaged.

## Acknowledgement

The authors acknowledge Crispin Hetherington and L. Tracy Yu for their technical assistance and insights. This work was funded in part by the National Science Foundation (CHE 2203937), the National Science Foundation Graduate Research Fellowship (Grant No. 1842494), the Welch Foundation (C-2141), the Swedish Research Council (2020-04633) and the NIH (GM122510). This work benefited from using the SasView application, originally developed under NSF award DMR-0520547. SasView contains code developed with funding from the European Union’s Horizon 2020 research and innovation program under the SINE2020 project, grant agreement No 654000. This work was partly done using the Shared Equipment Authority resources at Rice University.

## Supporting Information Available

Full peptide characterization, including UPLC, mass spectrometer data, CD melt curves, micrographs, models, and tables for mass balance.

## Author Contributions

Carson C. Cole: conceptualization, methodology, investigation, writing, data curation. Mark A.B. Kreutzberger: methodology, investigation, data curation, writing. Kevin Klein: methodology, investigation, data curation, writing. Kiana A. Cahue: investigation, data curation. Brett H. Pogostin: investigation, data curation. Adam C. Farsheed: investigation, data curation. Joseph W.R. Swain: investigation, data curation. Thi H. Bui: investigation, data curation. Arghadip Dey: investigation, data curation. Jonathan Makhoul: investigation, data curation. Marija Dubackic: investigation, data curation. Antara Pal: data curation. Ulf Olsson: conceptualization, funding acquisition. Andela Šarić: conceptualization, funding acquisition. Edward H. Egelman: supervision, conceptualization, funding acquisition. Jeffrey D. Hartgerink: conceptualization, editing, writing, supervision, funding acquisition.

## Conflicts of interest

The authors, J.D.H. and C.C.C. have filed a patent application based on the findings of this research

## Methods Continued

### Peptide Synthesis and Preparation

Peptides were manually synthesized using standard Fmoc solid phase peptide synthesis on a pre-loaded Gly-Wang resin. Fmoc-protected amino acids and resin were purchased from EMD Chemicals, and 2-(1H-7-azabenzotriazol-1yl)-1,1,3,3-tetramethyl uranium hexafluorophosphate methanaminium (HATU) was purchased from P3Bio. All chemicals not otherwise specified were purchased from Sigma-Aldrich. A mixture of 20% v/v piperidine mixture in dimethylformamide (DMF) was used for deprotecting steps. Coupling was performed using HATU and N,N-Diisopropylethylamine (DiEA) in DMF at 1:4:4:6 (resin/amino acid/HATU/DiEA, respectively). Acetylation of the N-terminus was performed twice with an excess of acetic anhydride and DiEA in dichloromethane. Cleavage was performed with 7.5% (v/v) scavengers (triisopropylsilane, MQ H_2_O, and anisole) in trifluoroacetic acid (TFA).

Excess TFA was removed after cleavage from the resin by a nitrogen flow. The crude peptide was triturated twice with cold diethyl ether. Crude peptides were dissolved in H_2_O to a 15 mg/mL concentration. This was sonicated and then filtered with a 0.3-micron syringe filter before purification by reverse-phase high-pressure liquid chromatography (HPLC) with water and MeCN with 0.05% TFA at a gradient of 0.85% per minute on a Waters XC18 column. Samples were roto-evaporated to remove MeCN and then lyophilized. Liquid chromatography mass spectrometry (LC-MS) was performed on an Agilent LC-MSD XT (Agilent Technologies, CA, USA) outfitted with a Pursuit 5 Agilent diphenyl column with the dimensions of 150 x 2.0 mm.

### UV-Vis Mass Balance

Samples were heated at 85°C for 30 minutes on heatblock to ensure all precipitates were dissolved. Absorbance values were measured at 280 nm using a Thermo Scientific™ NanoDrop™ 2000 Spectrophotometer. The measurements on day 0 were taken immediately after heating and stored at 4 ° C. On day 7, samples were centrifuged at 10000 rpm for 2 minutes, and the supernatant was measured. The percentage of insolubility was calculated using the following equation: ((1 – Abs*_Day_*_7_)/ Abs*_Day_*_0_) x 100=% Insoluble.

### Small Angle X-ray Scattering

Small-angle X-ray Scattering (SAXS) measurements were performed on the CoSAXS beam line, MAX IV, Sweden. Experiments were performed using an X-ray wavelength of 0.69 Å (18 KeV) and two sample-to-detector distances of 3.5 m and 0.5 m. Two-dimensional X-ray detection was achieved using an Eiger2 4M detector. The aqueous samples were filled into a 1.5 mm quartz capillary using a robotic autoloader. The samples were processed at a peptide concentration of 1 or 0.25% (w/w). Raw data were processed were processed using Python 3.11. Scattering data were modeled using SASview.

### Rheology

Rheological tests were performed on an AR-G2 rheometer (TA Instruments, New Castle, DE) with a 12 mm parallel plate. 70 µL of peptide was pipetted onto the stage, and the parallel plate was lowered to a gap of 550 µm. Any excess gel was scraped away before the gap was lowered to 500 µm, and oil was applied to prevent evaporation during testing.

**Figure S1:**
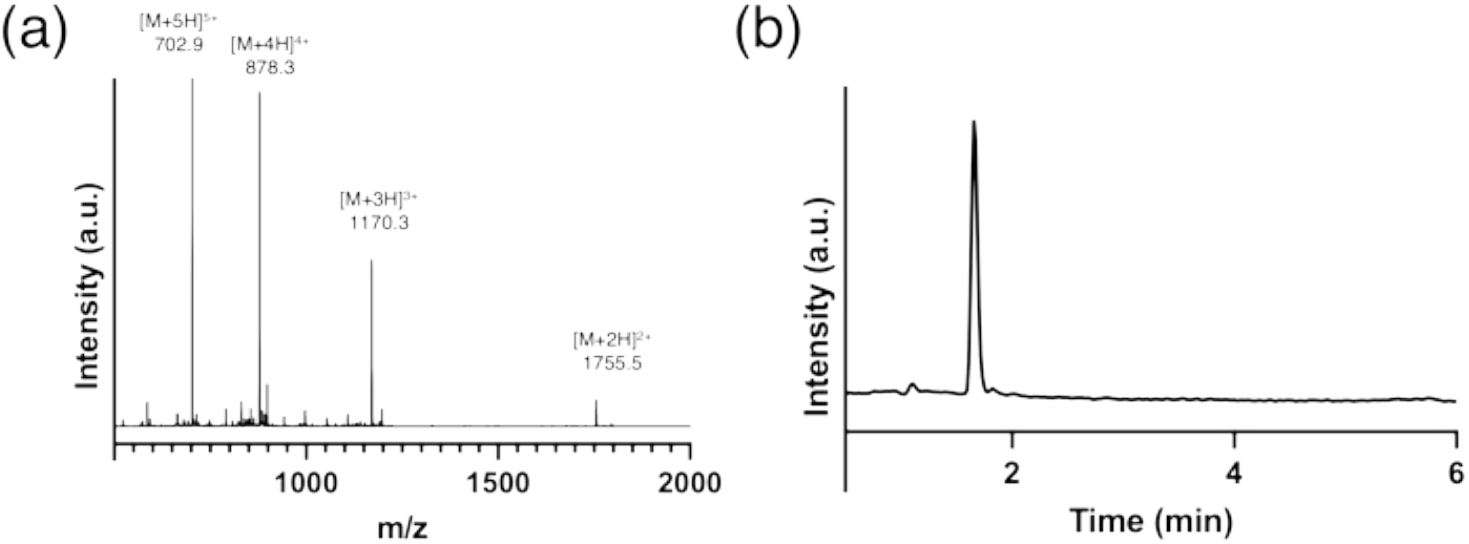
F_D2Y2_ Characterization: a) mass spectra (Expected: [M + H]^+^ = 3508.7; Observed: 3509.0) b) Pure liquid chromatography trace.

**Figure S2:**
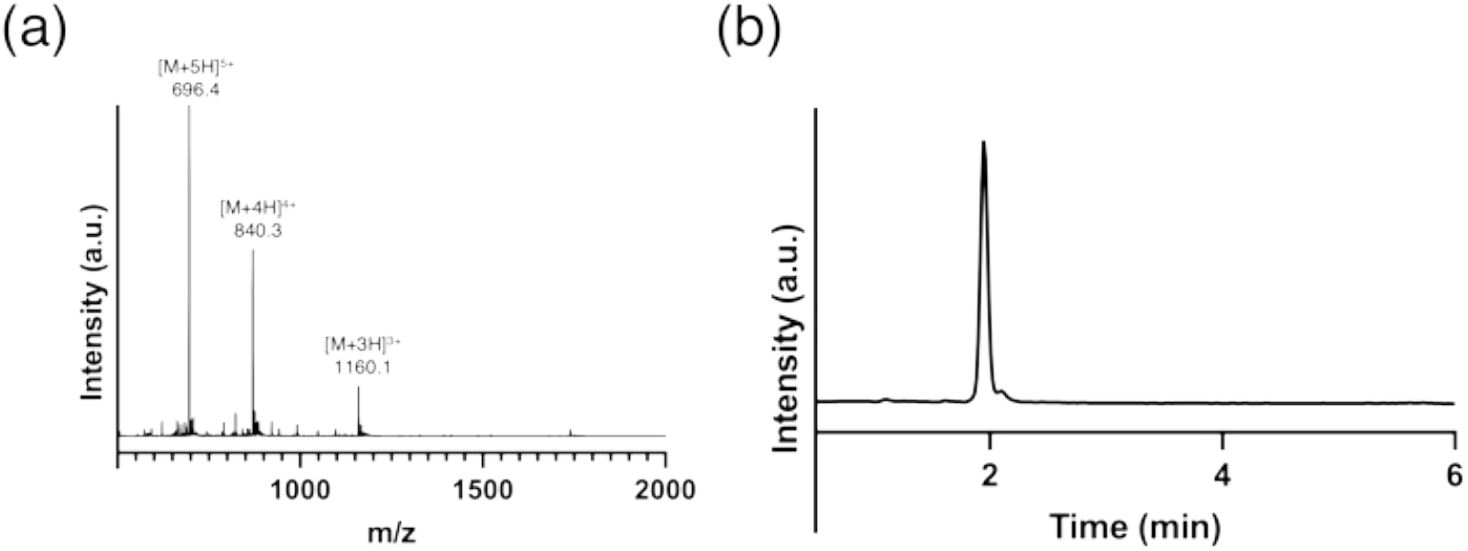
F_D2F2_ Characterization: a) mass spectra (Expected: [M + H]^+^ = 3476.7; Observed: 3477.3) b) Pure liquid chromatography trace.

**Figure S3:**
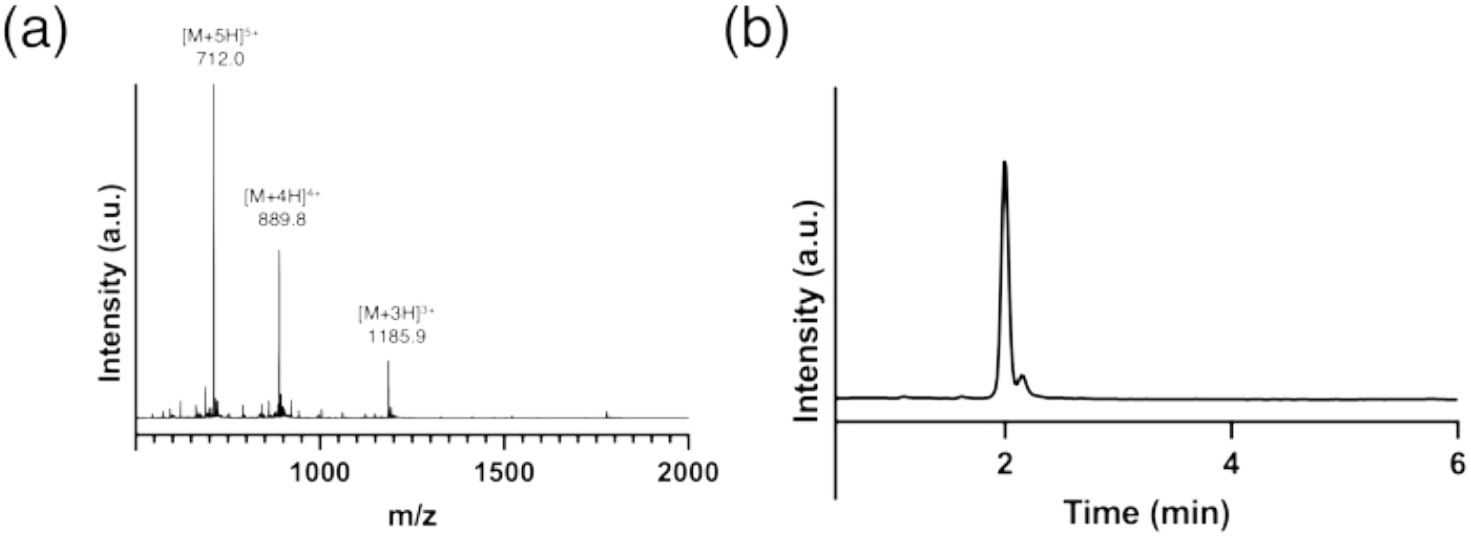
F_D2W2_ Characterization: a) mass spectra (Expected: [M + H]^+^ = 3554.7; Observed: 3554.7) b) Pure liquid chromatography trace.

**Figure S4:**
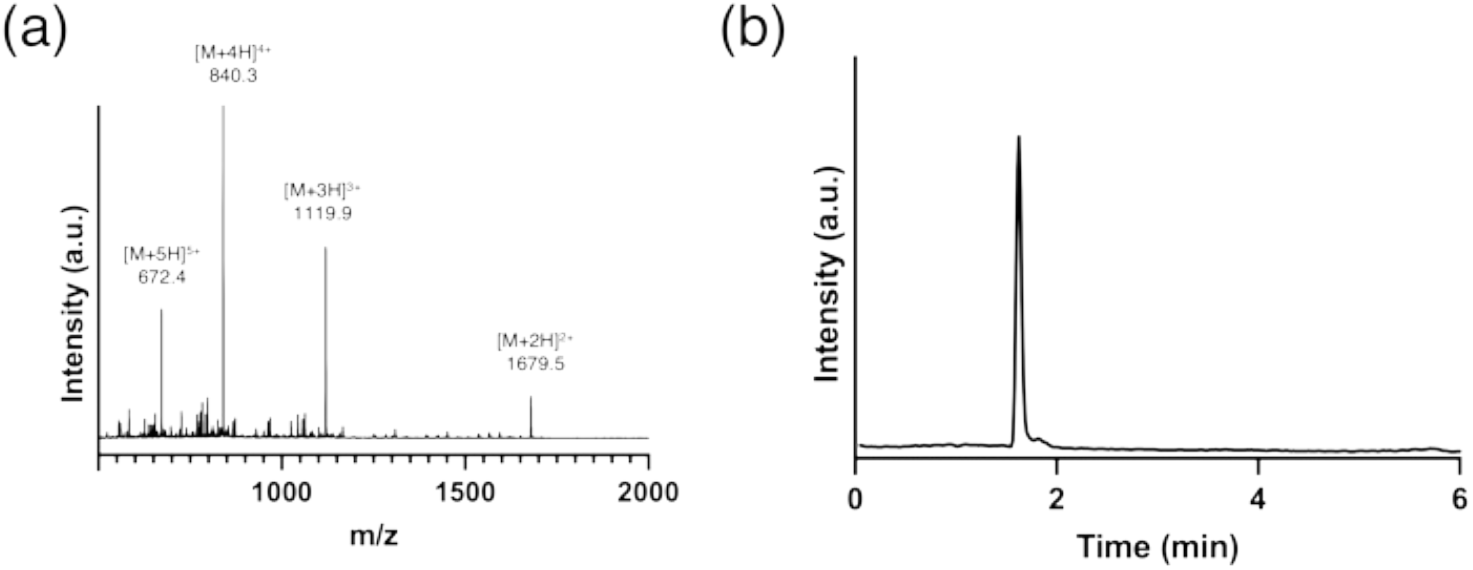
F_0_ Characterization: a) mass spectra (Expected: [M + H]^+^ = 3356.6; Observed: 3357.0) b) Pure liquid chromatography trace.

**Figure S5:**
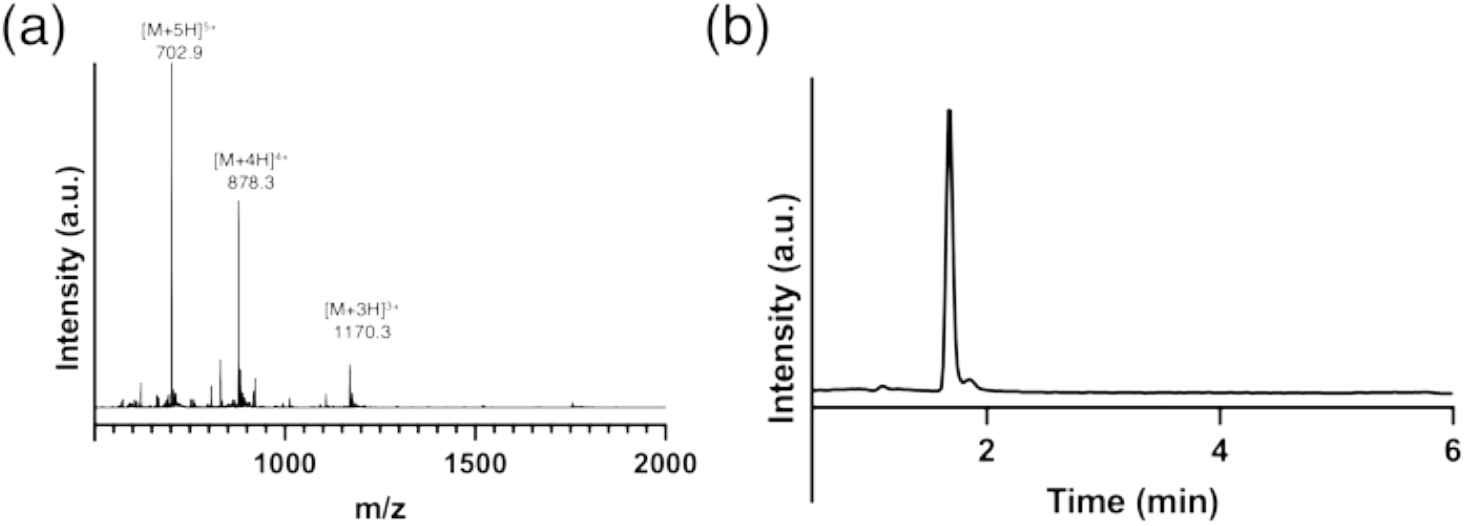
F_DY2_ Characterization: a) mass spectra (Expected: [M + H]^+^ = 3508.7; Observed: 3509.0) b) Pure liquid chromatography trace.

**Figure S6:**
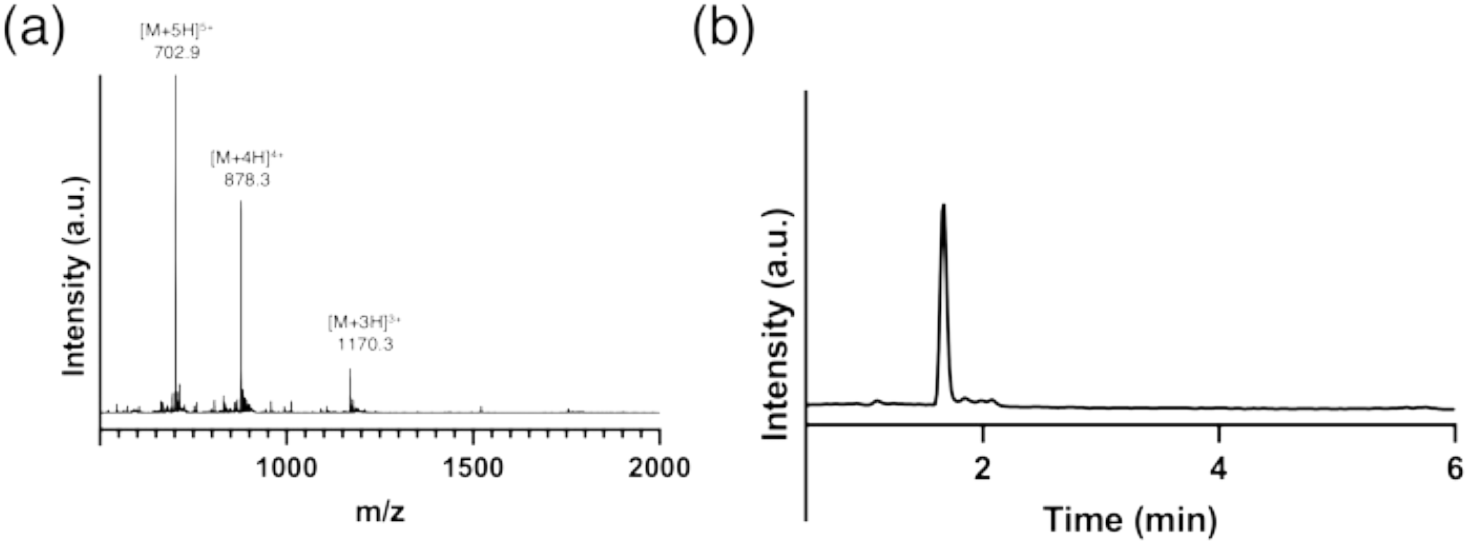
F_YD2_ Characterization: a) mass spectra (Expected: [M + H]^+^ = 3508.7; Observed: 3509.0) b) Pure liquid chromatography trace.

**Figure S7:**
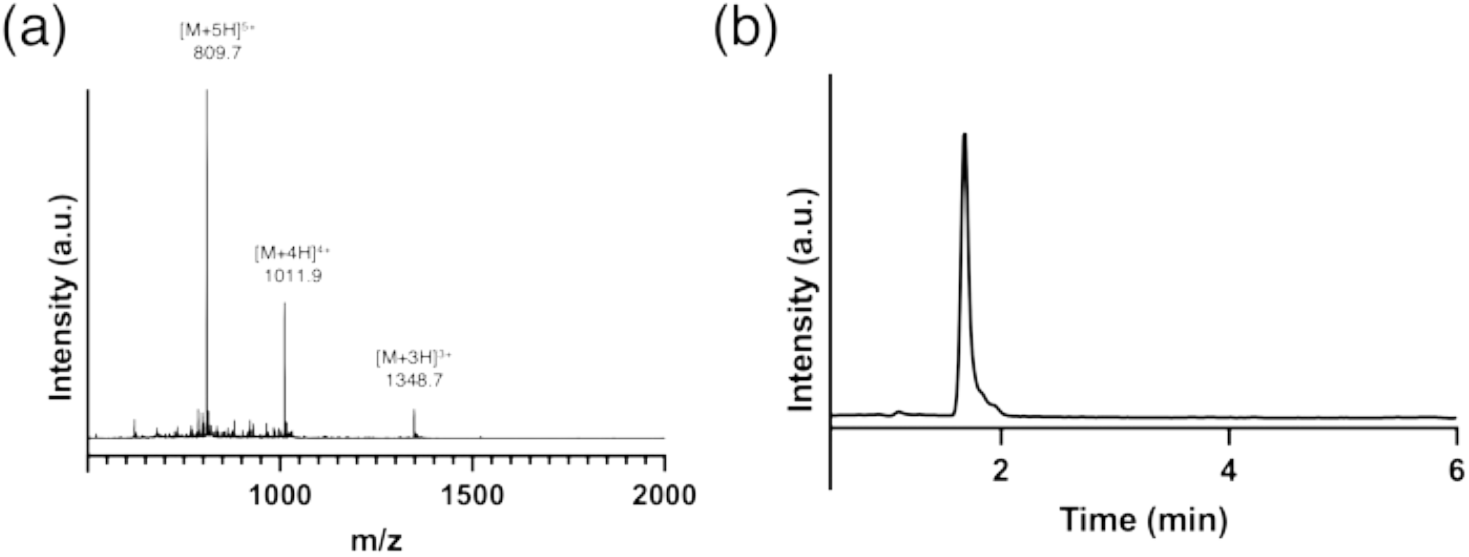
F_D2Y2_*_−_*_T_ Characterization: a) mass spectra (Expected: [M + H]^+^ = 4042.9; Observed: 4044.3) b) Pure liquid chromatography trace.

**Figure S8:**
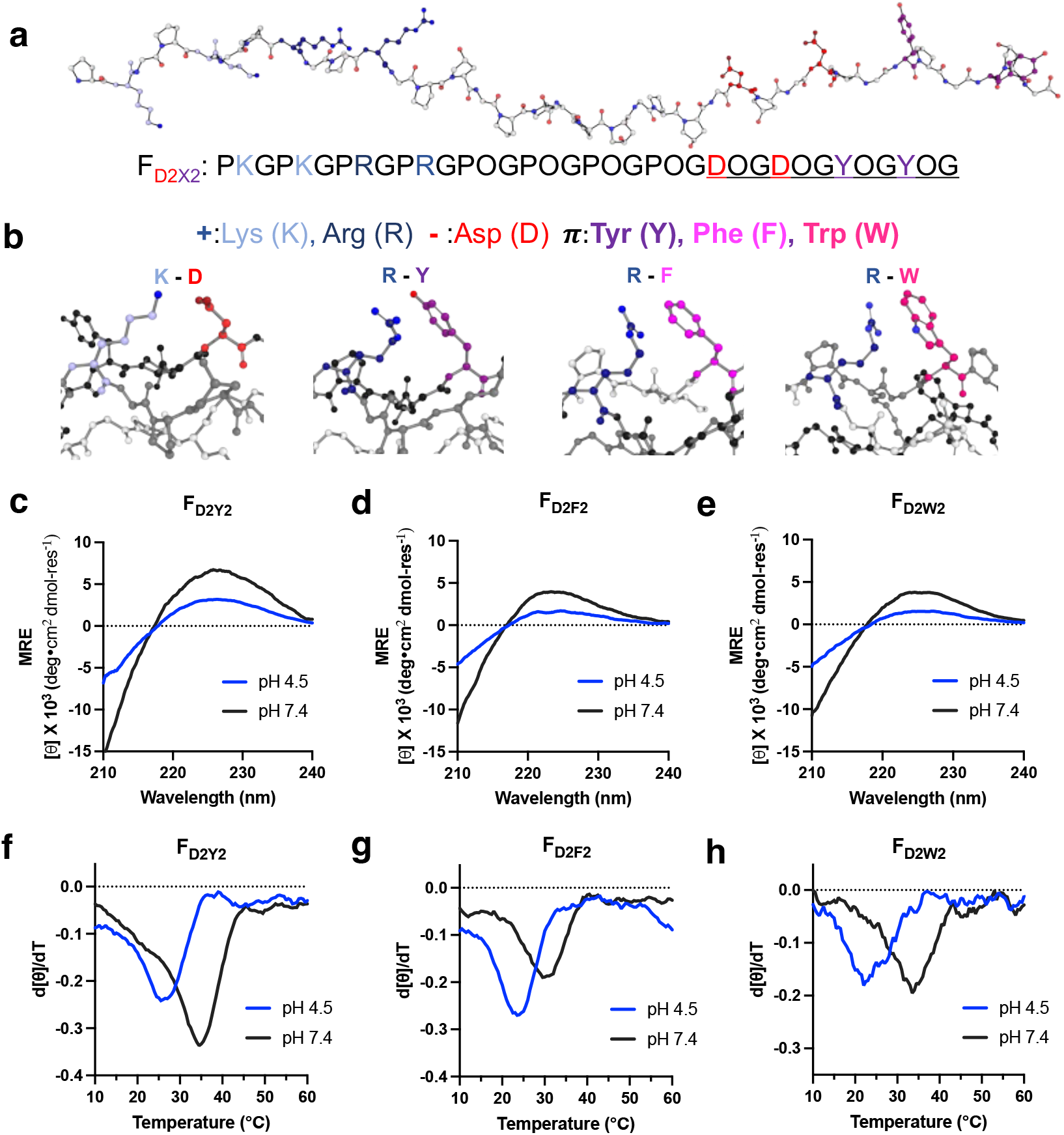
Circular dichroism of cation-*π* pairwise interaction screens. (a) A polyproline type II strand of F_D2Y2_ rendered with AlphaFold3.^1^ The underlined portion of the anionic and aromatic region corresponds to the nomenclature of the peptide. (b) Rendering of axial, intrahelical pairwise interactions, from left to right: lysine-aspartate charge pair, arginine-tyrosine, arginine-phenylalanine, and arginine-tryptophan cation-*π* interactions. Side chain conforms were optimized using DLPacker and using PDB: 8TW0.^2,3^ (c) Circular dichroism (CD) wavelength scans for phenylalanine-substituted F_D2Y2_ (d) CD scans for tryptophan-substituted F_D2F2_ (e) CD scans for tyrosine-substituted F_D2W2_ (f) CD thermal unfolding curves for phenylalanine-substituted F_D2Y2_ (g) CD thermal unfolding curves for tryptophan-substituted F_D2F2_ (h) CD thermal unfolding curves for tyrosine-substituted F_D2W2_. All CD spectra and melts were taken at 0.3 mM in MQ H_2_O.

**Figure S9:**
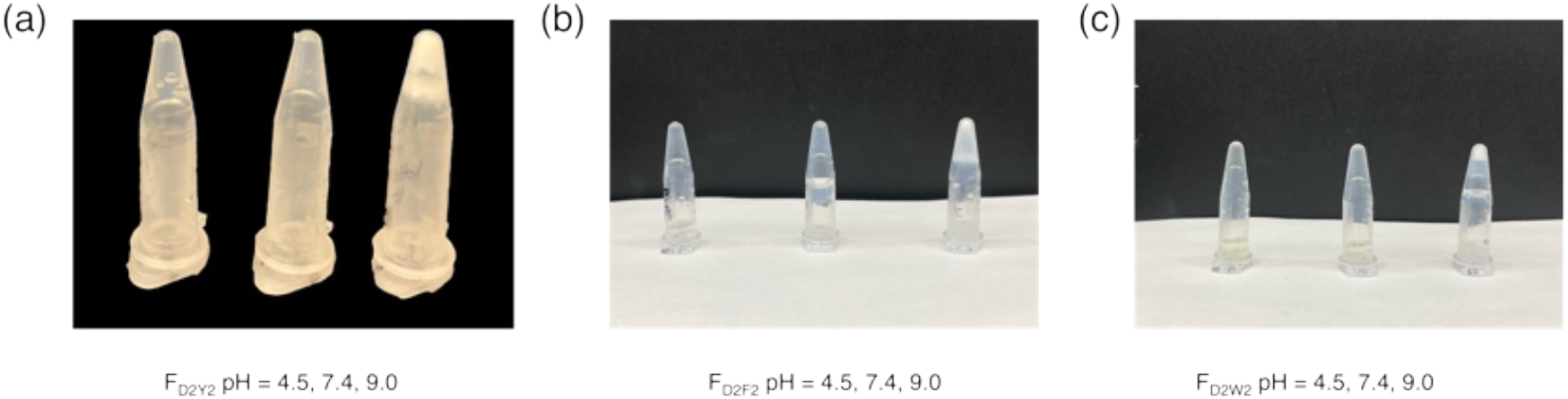
Inverted Eppendorf Tubes of D2X2 substituted CMPs at 2% w/v in MQ H2O. (a) FD2Y2 inverted eppendorf tubes. The pH 4.5 solution is a hydrogel and retains bubbles. (b) FD2F2 inverted eppendorf tubes. (c)FD2W2 inverted eppendorf tubes. Peptide samples were prepared in pH 4.5, 7.4, and 9.0 in MQ H2O, respectively. Precipitation is observed as pairwise interactions strengthen at higher pH.

**Figure S10:**
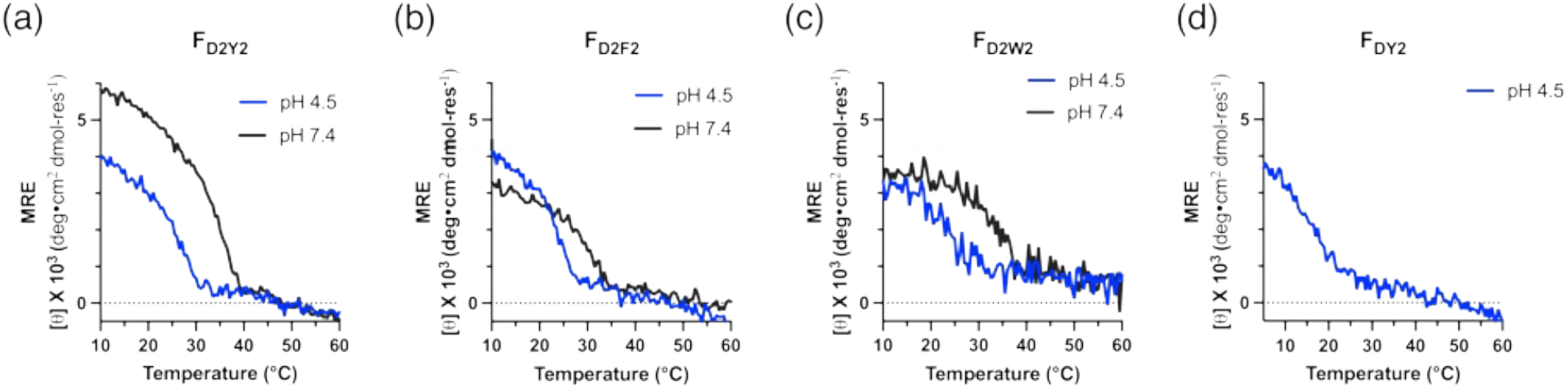
CD Thermal Melts of CMPs at pH 4.5 and 7.4 while being monitored at 225 nm (a) F_D2Y2_ thermal melt spectra (b)F_D2F2_ thermal melt spectra (c)F_D2W2_ thermal melt spectra. (d)F_DY2_ thermal melt spectra.

**Figure S11:**
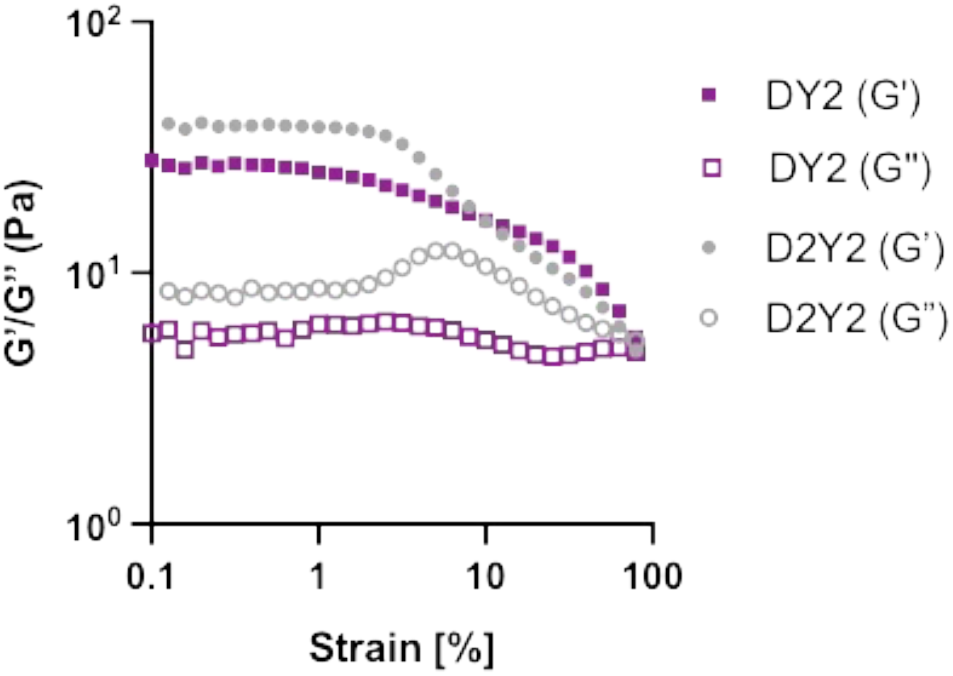
Rheology of F_D2Y2_ and F_DY2_ at pH 4.5 and 2 wt% in MQ H_2_O. Storage and loss moduli for each hydrogel. F_D2Y2_ has a storage and loss modulus of 39 Pa and 9 Pa, respectively. F_DY2_ has a storage and loss modulus of 32 Pa and 5 Pa, respectively.

**Figure S12:**
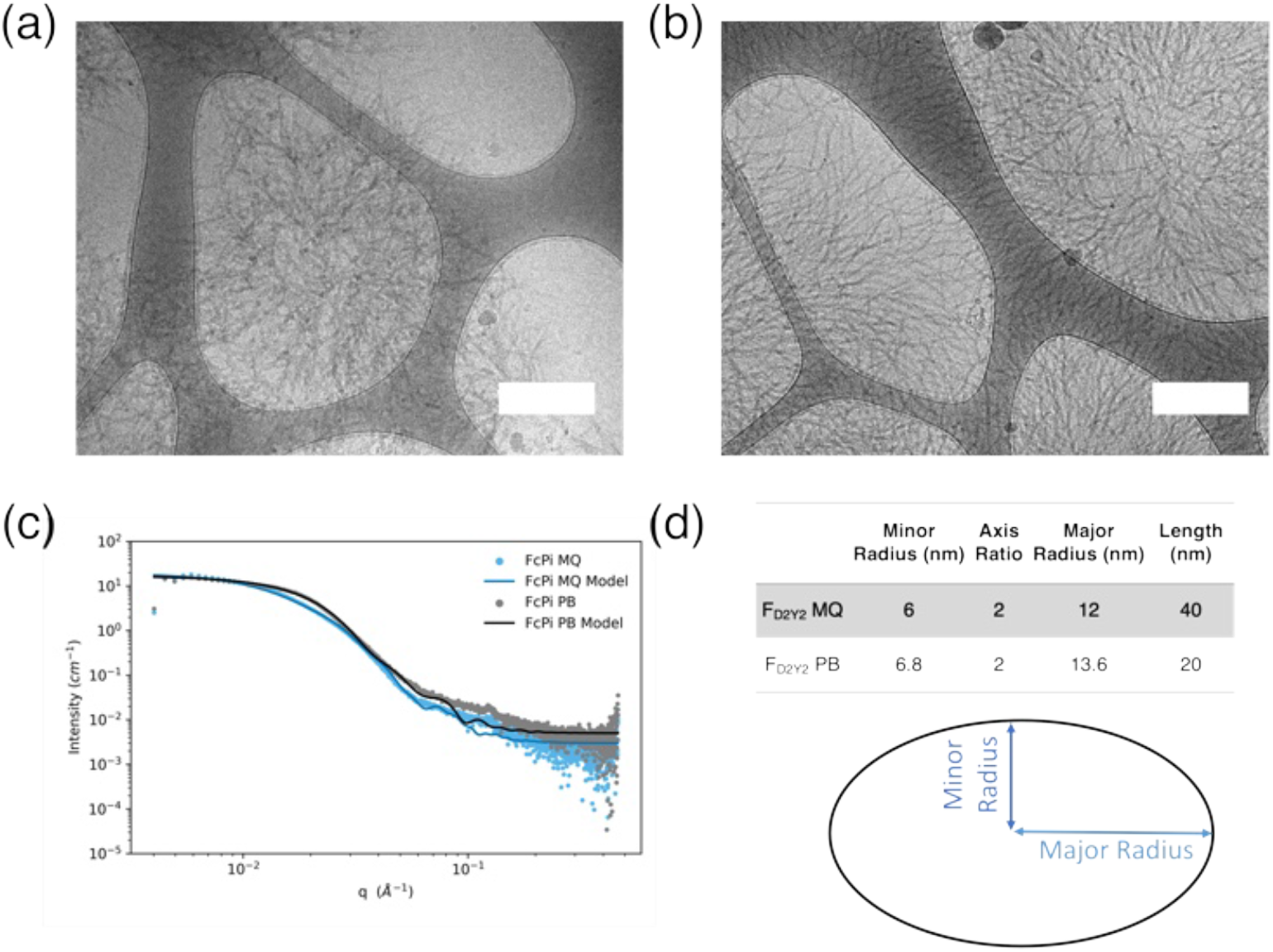
Phosphate buffer screen of F_D2Y2_ at pH 7.4 a) Cryo-EM of F_D2Y2_ in 10 mM phosphate buffer. Blank areas were present as there was a higher amount of association from the ion-containing buffer. b) Cryo-EM of F_D2Y2_ in H_2_O. Scale bars are 200 nm. (c) SAXS data for F_D2Y2_ in both H_2_O (MQ) and phosphate buffer. (PB) (d) Extrapolated dimensions of fibers in solution from SAXS data.

**Figure S13:**
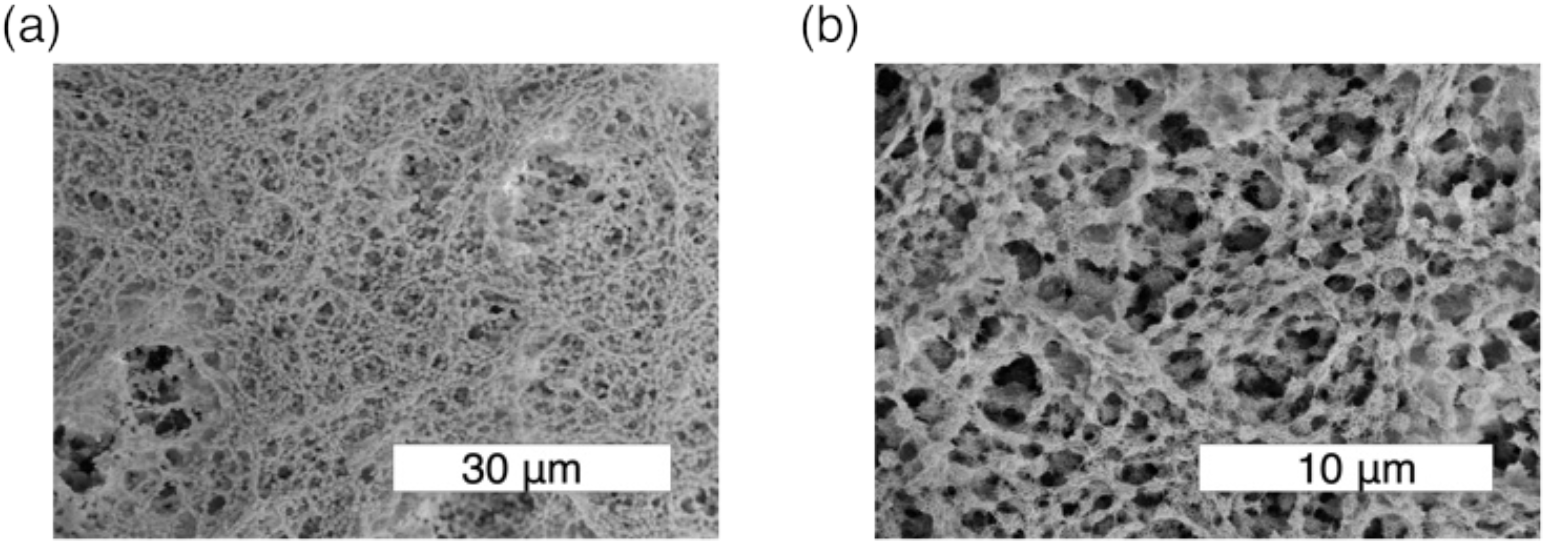
SEM of F_D2Y2_ at pH 7.4. (a) A zoomed-out SEM micrograph of 2 % (w/v) of F_D2Y2_. (b) A zoomed-in SEM micrograph of 2 % (w/v) of F_D2Y2_ showing distinct clustering of the collagen mimetic fibers.

**Figure S14:**
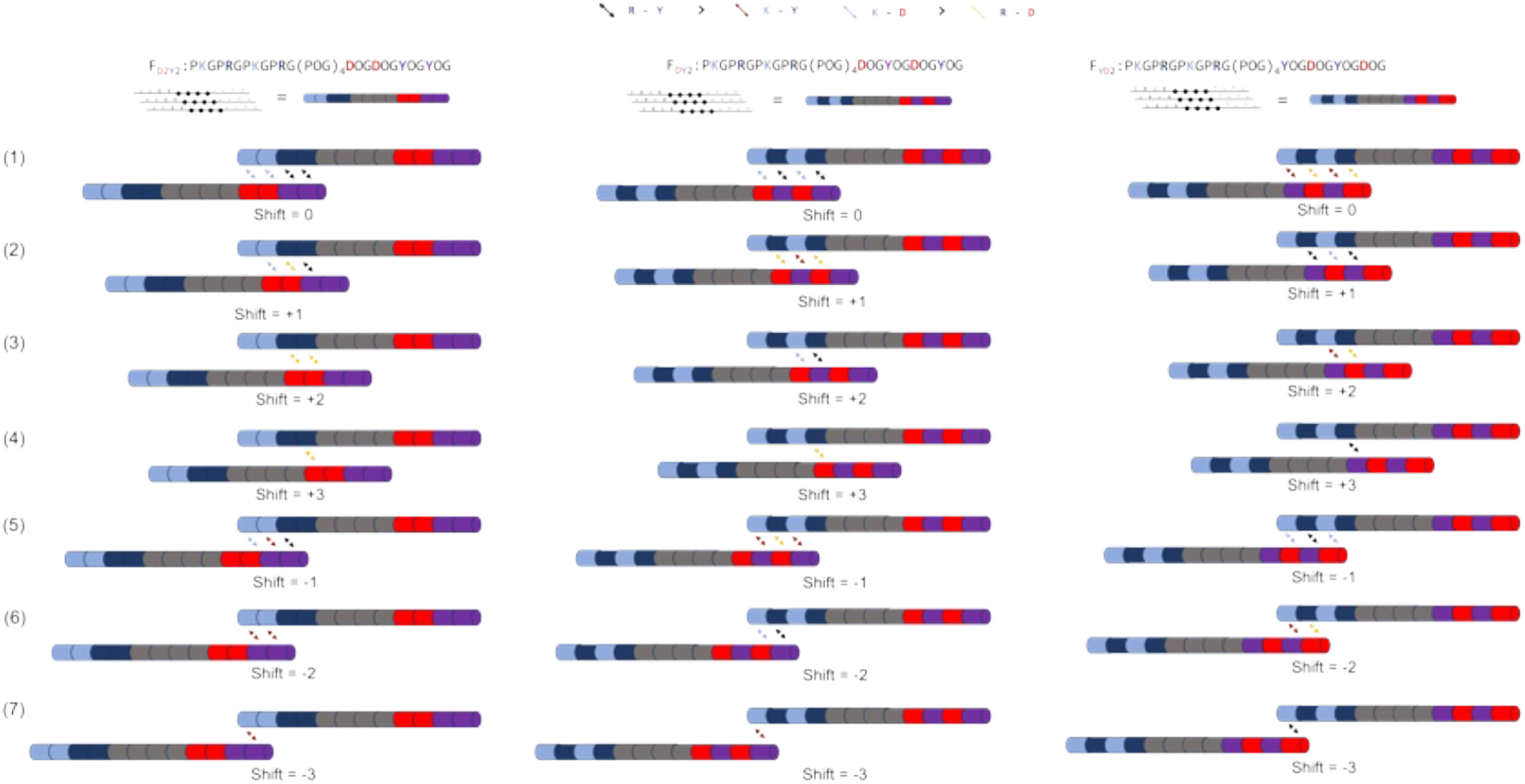
Possible parallel stagger comparisons of F_D2Y2_, F_DY2_ and F_YD2_ as triple helices shift past one another in space. Left: F_D2Y2_ staggers Center: F_DY2_ staggers Right: F_YD2_ staggers. Assuming electrostatic charge pairs are also involved.

**Figure S15:**
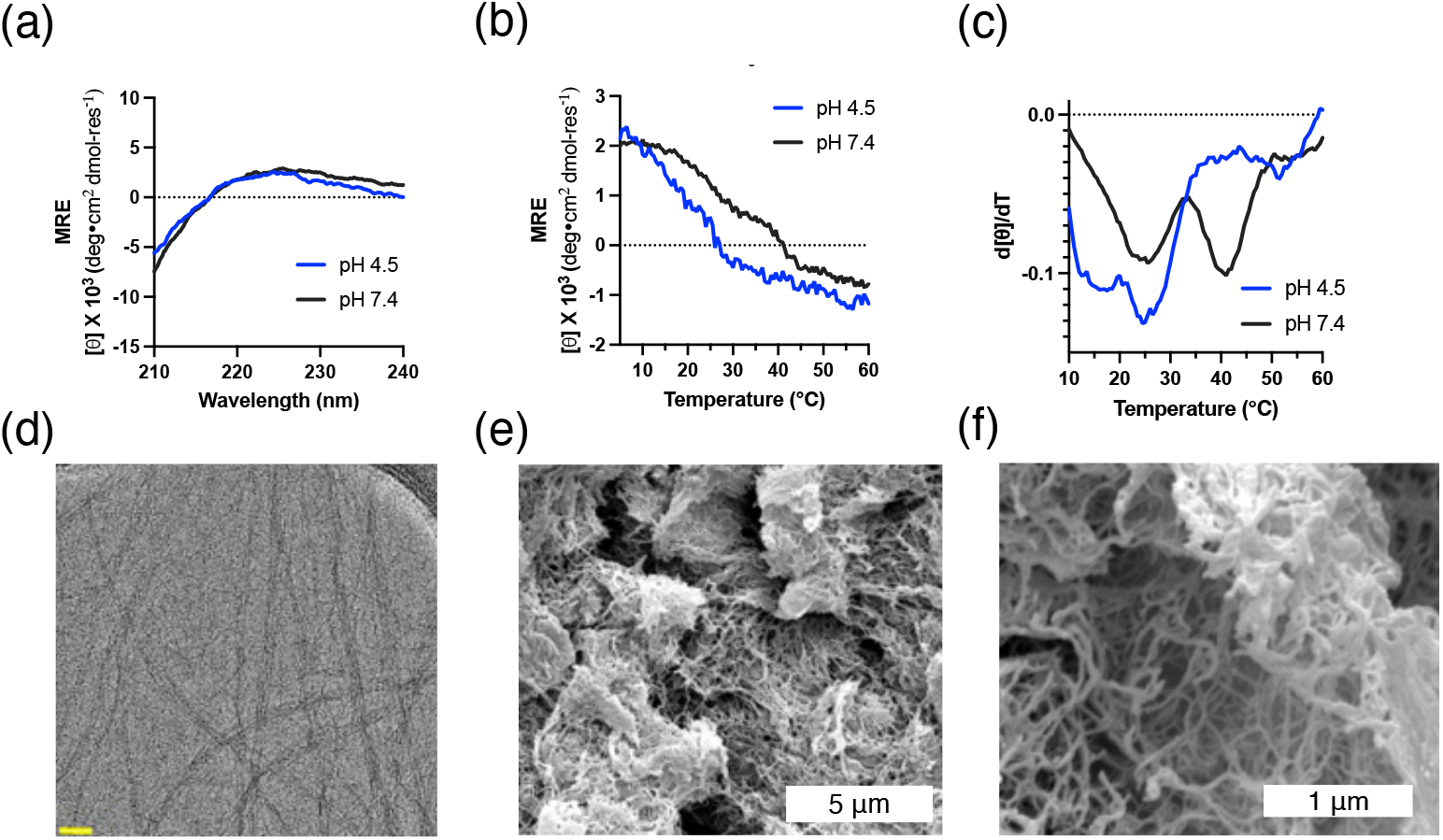
Circular dichroism and electron microscopy of F0. (a) CD spectra of F_0_. (b) CD thermal melt curves of F_0_ monitored at 225 nm. (c) CD first-order derivative of F_0_ thermal melt curves. (d) Cryo-EM pf F_0_ shows multiple fibers associating at pH 7.4. The scale bar is 40 nm. (e) SEM of F_0_ shows a fibrous matrix. (f) Zoomed in SEM micrograph of F_0_’s fibrous matrix. Peptides were dissolved at 1.0 % w/v in 10 mM phosphate buffer and prepared as previously reported.^4^

**Figure S16:**
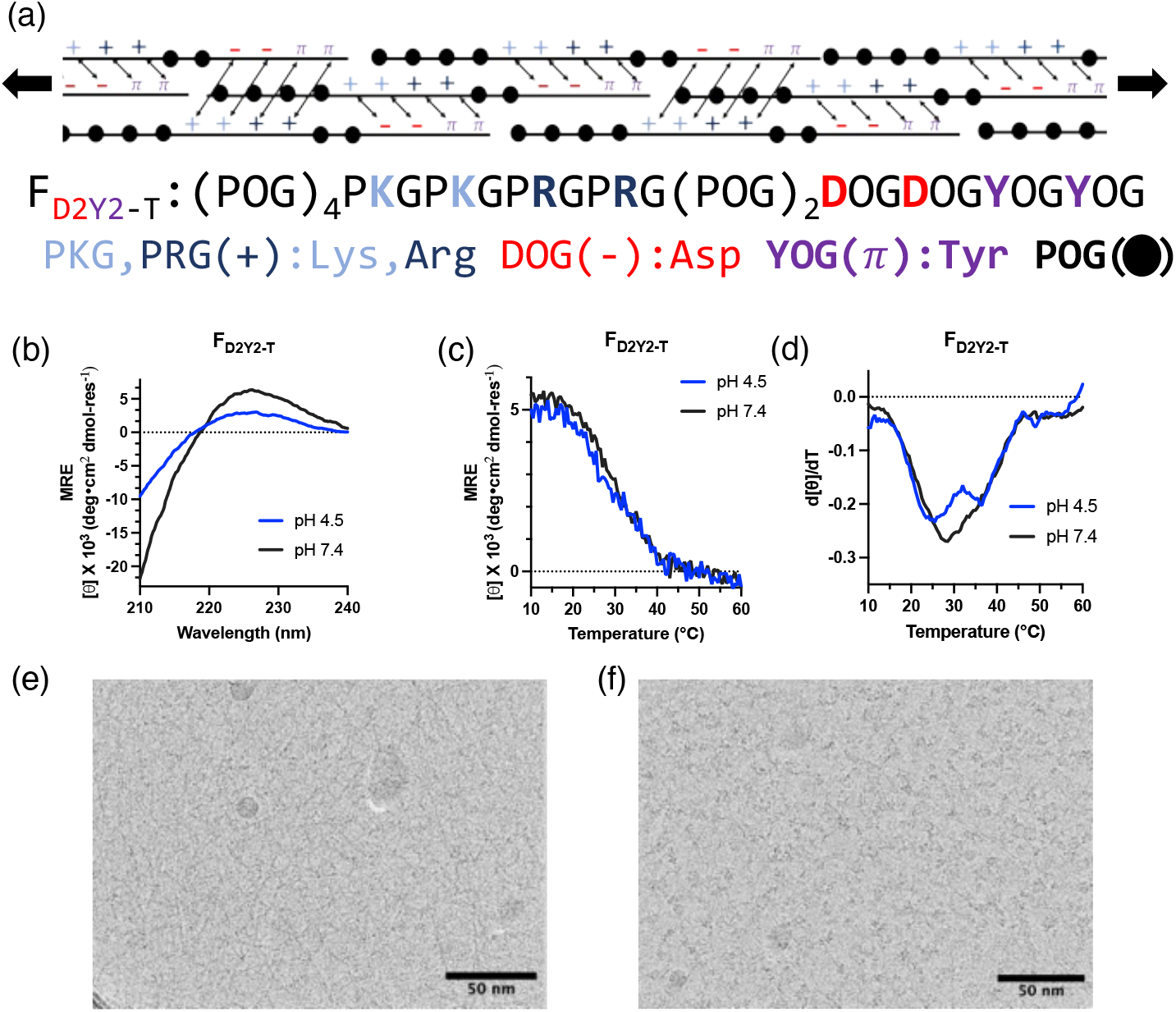
Circular dichroism and electron microscopy of F_D2Y2T_ . (a) The proposed assembly mechanism of F_D2Y2T_ could result in satisfied intrahelical interactions. (a) CD spectra of F_D2Y2T_ from pH 4.5 and 7.4. (b) CD thermal melt curves of F_D2Y2T_ monitored at 225 nm. (c) CD first-order derivative of F_D2Y2T_ thermal melt curves. (d) Cryo-EM pf F_D2Y2T_ shows multiple fibers at pH 4.5. The scale bar is 50 nm. (e) Commonly observed aggregates of F_D2Y2T_ in Cryo-EM at pH 4.5. The scale bar is 50 nm. Samples were prepared from a 2.0 % w/v solution in MQ H_2_O.

**Figure S17:**
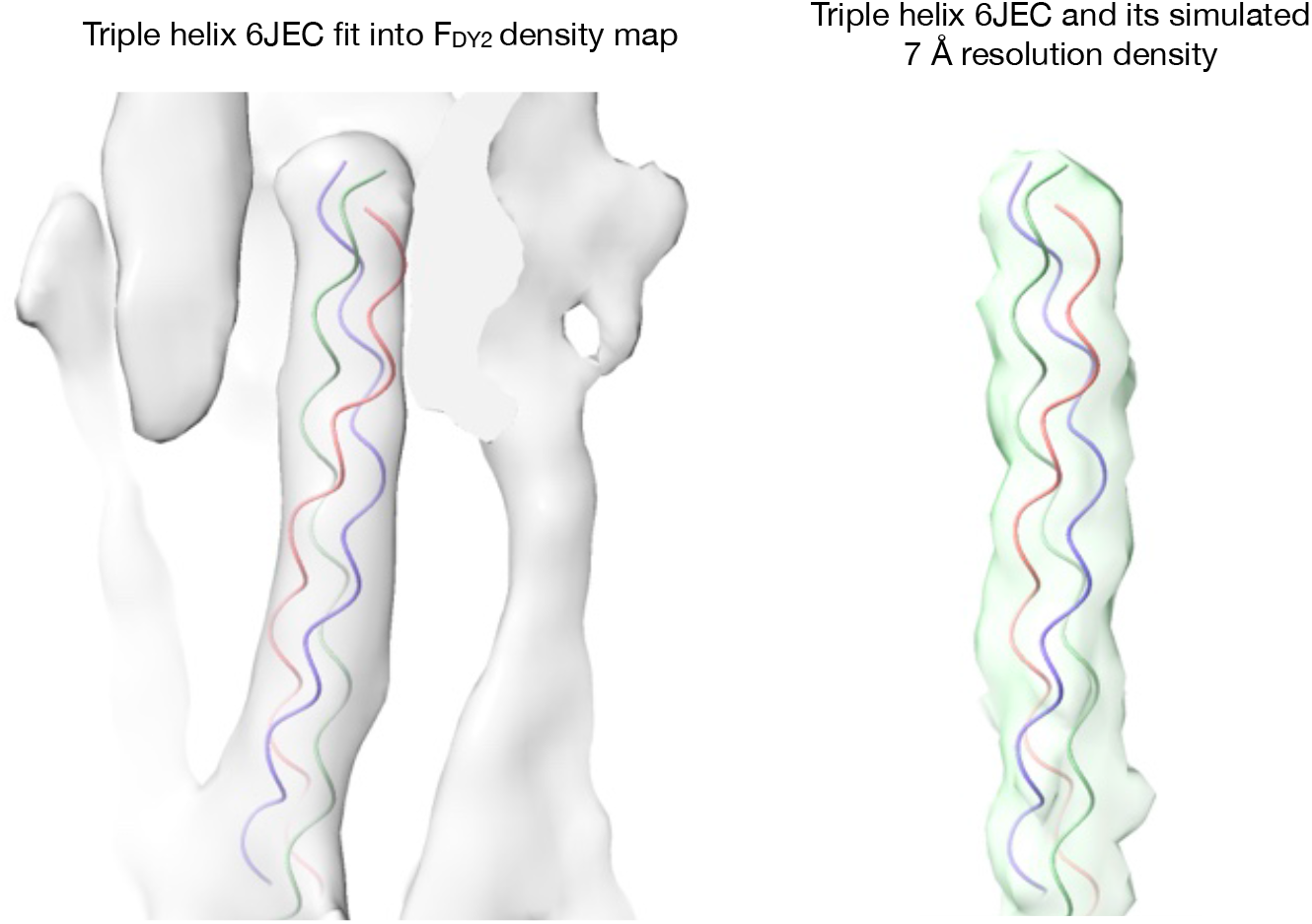
3D resolution comparison of F_DY2_ and PDB 6JEC. The left image shows a ribbon representation of a triple helix (6JEC) fit into the FDY2 density map (grey). The right image shows the model 6JEC and its a simulated density map at 7 Å resolution (green) generated from the molmap command in UCSF ChimeraX.

**Figure S18:**
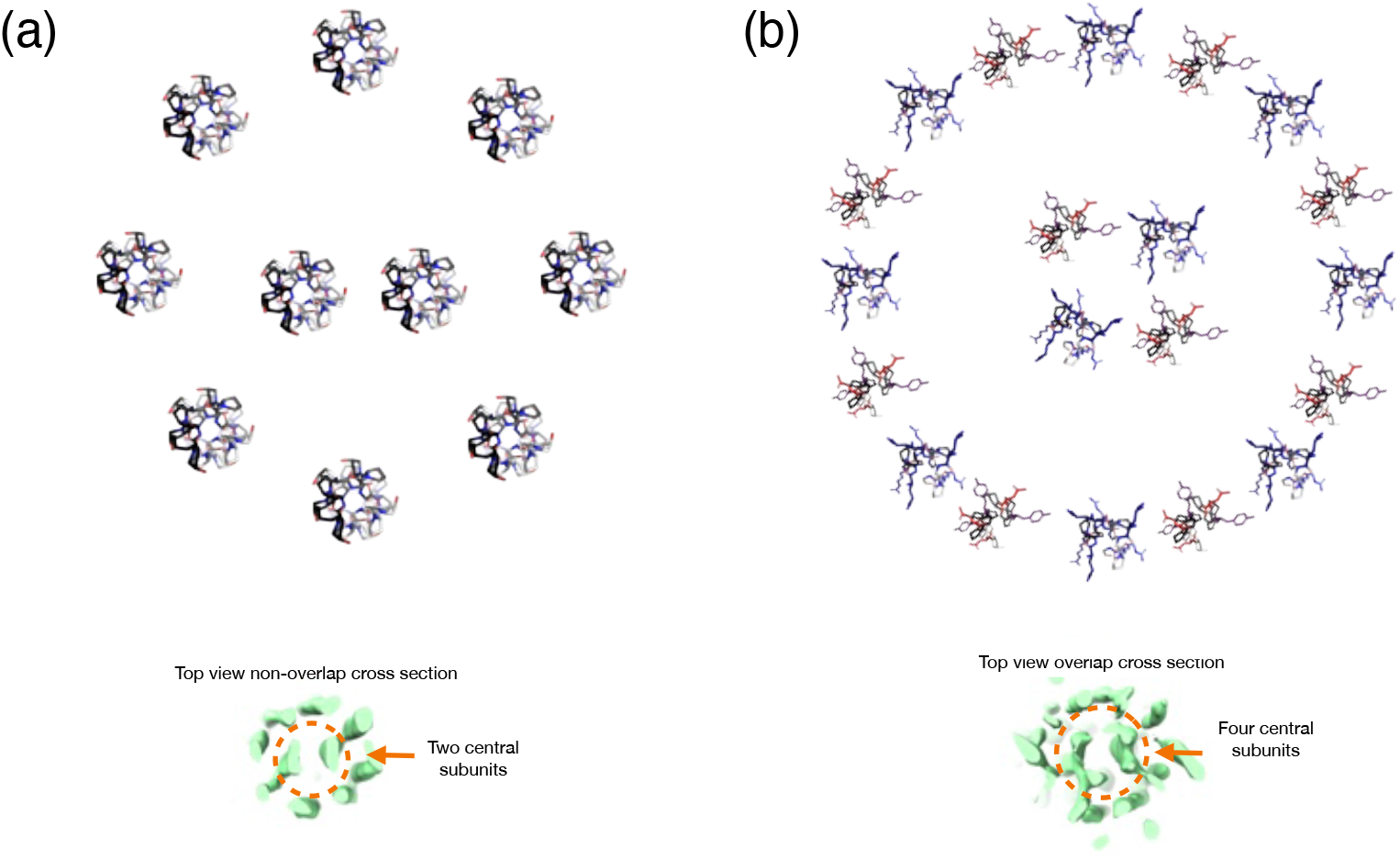
Possible triple helix orientation to explain 8+2 stacking of F_DY2_ (a) Hydrophobic slice of the central POG region in F_DY2_ compared with a slice from the experimental Cryo-EM data. This correlates to the lighter band of the D-band observed in Figure 5 of the main manuscript. (b) Compared with a slice from the experimental Cryo-EM data, the overlap slice of the terminal regions in F_DY2_. This proposes a theoretical 16+4 packing in the overlap region corresponding to the dark portion observed in the D-banding. The triple helical backbone was prepared using AlphaFold3.^1^

**Figure S19:**
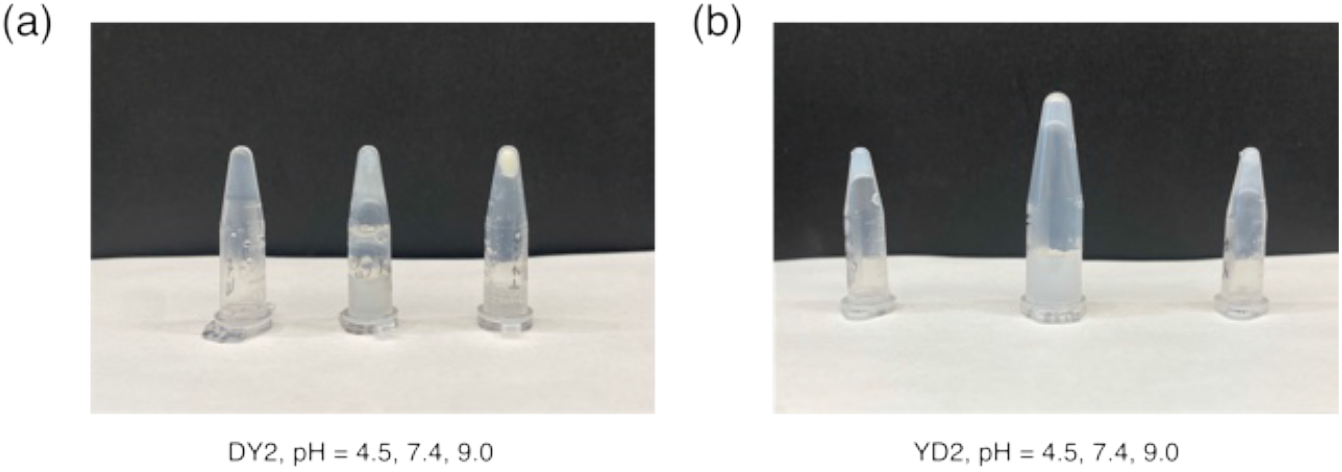
Eppendorf tubes of F_DY2_ and F_YD2_ at various pH values. (a) F_DY2_ samples prepared 2.0 wt % in MQ H_2_O. (b) F_YD2_ samples prepared 2.0 wt % in MQ H_2_O.

**Figure S20:**
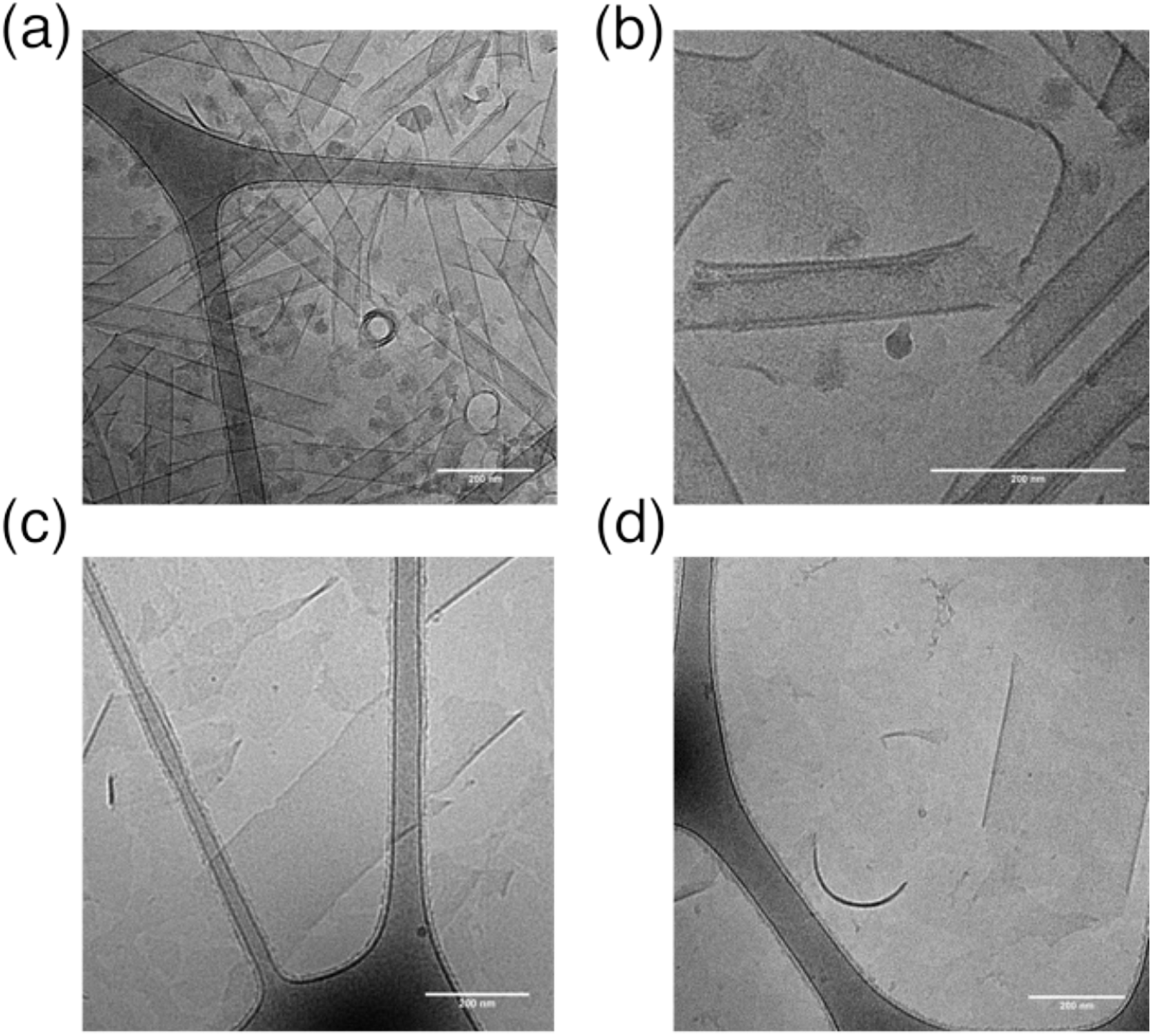
Further Cryo EM of F_DY2_ and F_YD2_ in MQ H_2_O (a) Cryo-EM of F_DY2_ nanotube micrographs show the cross-section of tubes. (b) F_DY2_ nanotube appear to be multi-lamellar. (c) F_YD2_ nanosheets appear to be broken at the center of the micrograph.(d) A cross-section of the nanosheet shows a curling of the sides. Precipitation is observed in all F_YD2_ samples. F_DY2_ is a hydrogel at pH 4.5 and solvated. All scale bars are 200 nm.

**Figure S21:**
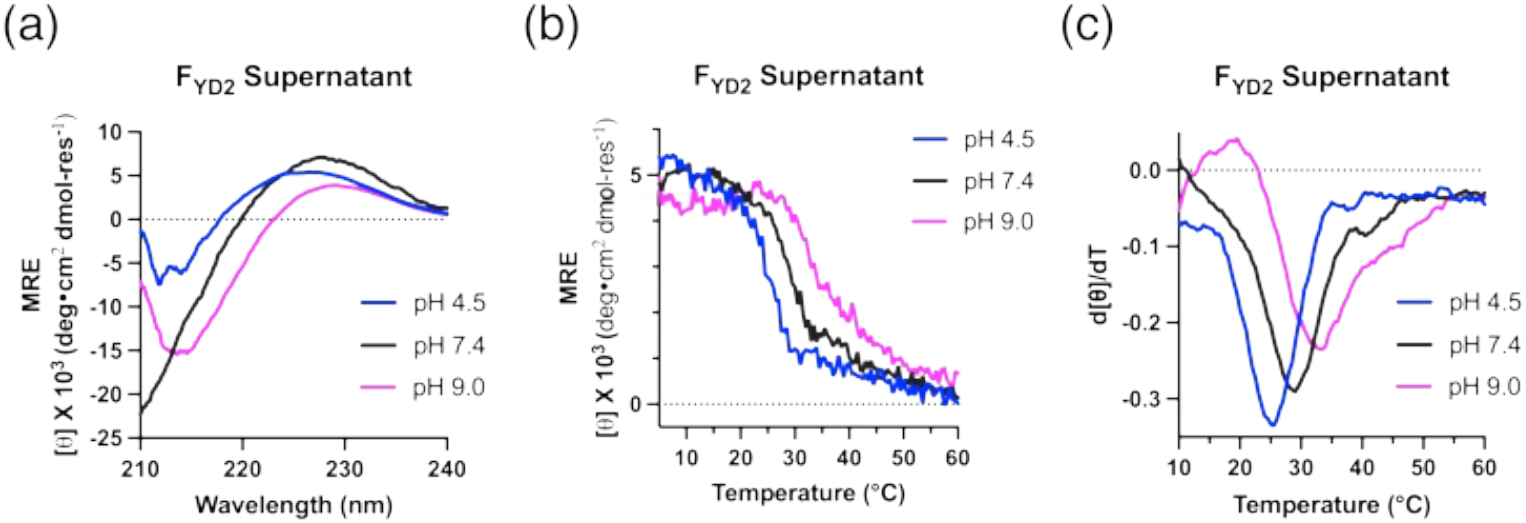
Circular dichroism characterization of F_YD2_ supernatant at various pH values. (a) CD scans of supernatant of F_YD2_ (b) CD thermal melt curves of F_YD2_ monitored at *∼*225 nm. (c) First derivative of the thermal melt curves. All samples were prepared at specified pH and collected at 0.3 mM in MQ H_2_O.

**Figure S22:**
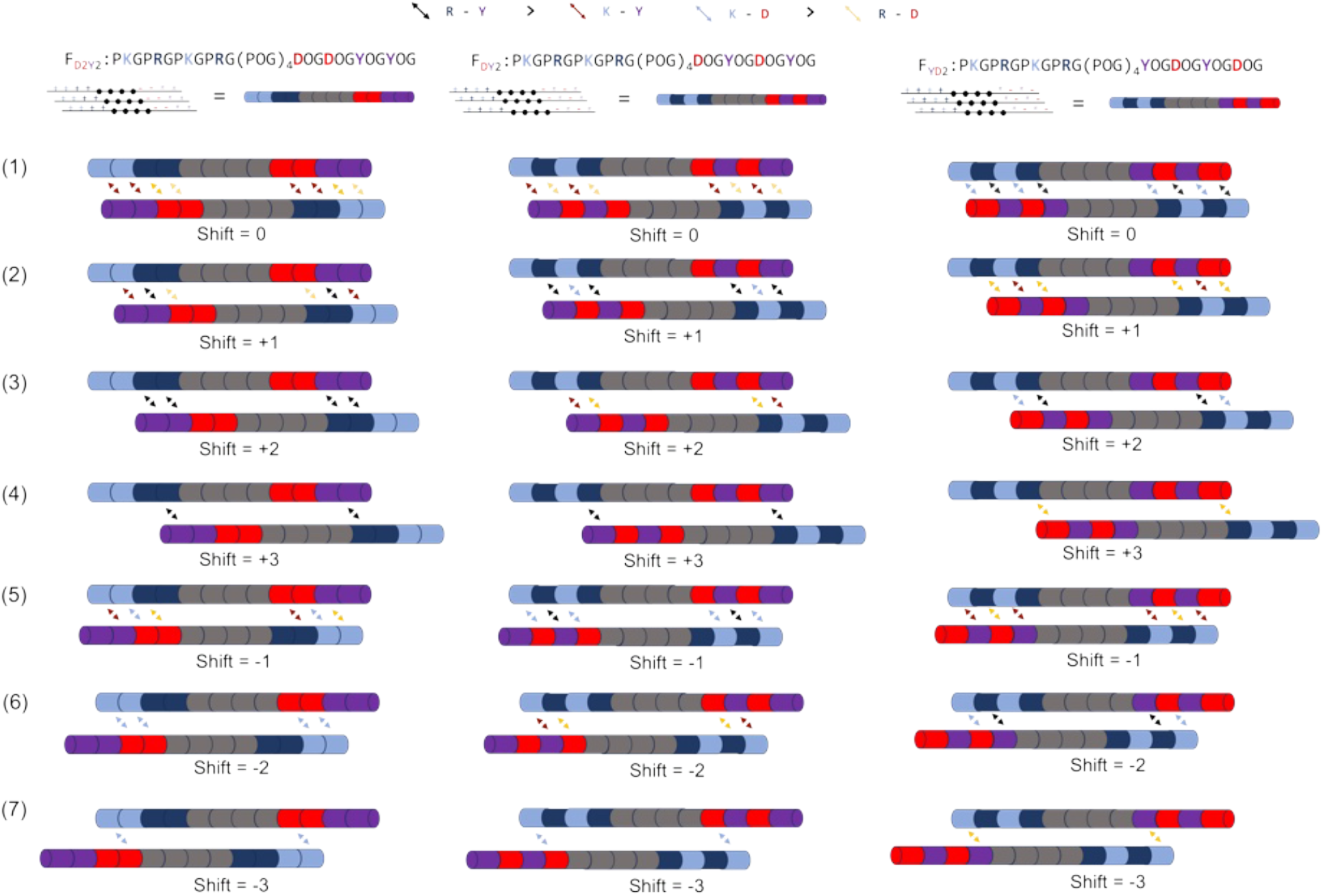
Possible parallel stagger comparisons of F_D2Y2_, F_DY2_ and F_YD2_ Left: F_D2Y2_ staggers between strands show that the energy gap is in comparison with stagger (1) and staggers (2) and (5). Right: F_DY2_ staggers between strands show that the energy gap is in comparison with stagger (1) and staggers (2), (3) and (6), but the energy difference between the most stable (1) is greater than that for F_D2Y2_ and its competing staggers. Relative pairwise interactions were deconvolved at pH 7.4.

**Table S1:** Peptide Thermal Stability as Determined by Circular Dichroism

| Sample | Sequence | pH 4.5 | pH 7.4 | $\Delta T$ |
| --- | --- | --- | --- | --- |
| $F_{D2F2}$ | (PKG)<sub>2</sub>(PRG)<sub>2</sub>(POG)<sub>4</sub>(DOG)<sub>2</sub>(FOG)<sub>2</sub> | 24.0 °C | 30.0 °C | 6.0 °C |
| $F_{D2W2}$ | (PKG)<sub>2</sub>(PRG)<sub>2</sub>(POG)<sub>4</sub>(DOG)<sub>2</sub>(WOG)<sub>2</sub> | 22.0 °C | 34.0 °C | 12.0 °C |
| $F_{D2Y2}$ | (PKG)<sub>2</sub>(PRG)<sub>2</sub>(POG)<sub>4</sub>(DOG)<sub>2</sub>(YOG)<sub>2</sub> | 26.5 °C | 34.5 °C* | 8.0 °C |
| $F_{DY2}$ | (PKGPRG)<sub>2</sub>(POG)<sub>4</sub>(DOGYOG)<sub>2</sub> | 15.5 °C | —* | |
| $F_{YD2}$ | (PKGPRG)<sub>2</sub>(POG)<sub>4</sub>(YOGDOG)<sub>2</sub> | —* | —* | |
| $F_0$ | (PKG)<sub>4</sub>(POG)<sub>4</sub>(DOG)<sub>4</sub> | 25.0 °C | 41.0 °C | 16.0 °C |
| $F_{D2Y2-T}$ | (POG)<sub>4</sub>(PKG)<sub>2</sub>(PRG)<sub>2</sub>(POG)<sub>2</sub>(DOG)<sub>2</sub>(YOG)<sub>2</sub> | 25.5 °C | 29.0 °C | 3.5 °C |

Cations - Blue, Anions - Red, Aromatic - Violet; \*Precipitate observed. Reported thermal stability values ( $T_m$ ) are the minima of the first-order derivative curves. All sample concentrations were measured at 0.3 mM and pH-adjusted with HCl and NaOH in H<sub>2</sub>O.

**Table S2:** Mass balance of F_D2Y2_, F_DY2_ and F_YD2_ based on absorbance at 280 nm

|  | D2Y2 pH 7.4 | pH 9.0 | DY2 pH 7.4 | pH 9.0 | YD2 pH 4.5 | pH 7.4 | pH 9.0 |
| --- | --- | --- | --- | --- | --- | --- | --- |
| Day 0 Abs. | 1.69 | 2.17 | 1.67 | 2.22 | 1.69 | 1.85 | 2.86 |
| Day 7 Abs. | 1.38 | 0.35 | 0.53 | 0.43 | 1.24 | 0.90 | 0.38 |
| Insoluble % | 18.8% | 84.1% | 68.2% | 80.8% | 26.5% | 51.2% | 86.7% |

All peptides were measured from a 2 % w/v in H<sub>2</sub>O and taken from the same parent stock.
Absorbance values are arbitrary units. All samples taken at 4 °C.

**Table S3:** Fiber Forming Peptides Sequences and pH Dependency for Hydrogel Formation

| name | sequence | pI | pH 4.5 | pH 7.4 | pH 9.0 |
| --- | --- | --- | --- | --- | --- |
| F0 | (PKG)<sub>4</sub>(POG)<sub>4</sub>(DOG)<sub>4</sub> | 7.2 | – | ✓ | P |
| F<sub>D2F2</sub> | (PKG)<sub>2</sub>(PRG)<sub>2</sub>(POG)<sub>4</sub>(DOG)<sub>2</sub>(FOG)<sub>2</sub> | 10.8 | – | – | P |
| F<sub>D2W2</sub> | (PKG)<sub>2</sub>(PRG)<sub>2</sub>(POG)<sub>4</sub>(DOG)<sub>2</sub>(WOG)<sub>2</sub> | 10.8 | – | – | P |
| F<sub>D2Y2</sub> | (PKG)<sub>2</sub>(PRG)<sub>2</sub>(POG)<sub>4</sub>(DOG)<sub>2</sub>(YOG)<sub>2</sub> | 10.2 | ✓ | P | P |
| F<sub>(DY)2</sub> | (PKGPRG)<sub>2</sub>(POG)<sub>4</sub>(DOGYOG)<sub>2</sub> | 10.2 | ✓ | P | P |
| F<sub>(YD)2</sub> | (PKGPRG)<sub>2</sub>(POG)<sub>4</sub>(YOGDOG)<sub>2</sub> | 10.2 | P | P | P |
| F<sub>D2Y2-T</sub> | (POG)<sub>4</sub>(PKG)<sub>2</sub>(PRG)<sub>2</sub>(POG)<sub>2</sub>(DOG)<sub>2</sub>(YOG)<sub>2</sub> | 10.2 | – | – | – |

Cations - Blue, Anions - Red, Aromatic - Violet; P: denotes a precipitate formed in solution at 2 % w/v in H<sub>2</sub>O; A check-mark denotes the formation of a hydrogel; – indicates a clear solution; pI is the isoelectric point.

## Notes

### Competing Interest Statement

The authors have declared no competing interest.

